# Ancestry and Environmental Adaptation in North American Feral Cannabis

**DOI:** 10.64898/2026.02.05.704066

**Authors:** Anna Halpin-McCormick, Ademola Aina, Michael Kantar, Shelby Ellison

## Abstract

Understanding how different populations respond to environmental variation is fundamental to breeding climate resilient crops. In this study, we integrate three diverse georeferenced Cannabis datasets comprising North American feral populations and Eurasian samples (n=909) to resolve population structure, infer evolutionary relationships and quantify adaptive responses to climate across a broad environmental gradient. Phylogenetic analysis rooted to *Humulus lupulus* shows North American feral populations are more closely related to basal and hemp-type lineages than to drug-type or Iranian populations, a pattern supported by ancestry and PCA analyses. By combining these datasets, we capture a wider range of climatic variation and gain new insight into the adaptive potential of cannabis germplasm. Using environmental genomic selection (EGS), we identified nine bioclimatic traits with prediction accuracies exceeding 0.5 across the combined datasets (12,030 SNPs; 909 samples; training = 191). When analyzing the North American dataset alone, EGS revealed 14 traits with prediction accuracies greater than 0.85 (22,852 SNPs; 760 samples; training = 310). Genomic estimated adaptive values (GEAVs) revealed population specific climatic responses, particularly for precipitation related traits, with northeastern North American populations (Indiana and New York) showing signatures consistent with adaptation to wetter and cooler environments. Climate projections under a high emission scenario (SSP585) indicate that ∼34 % of sampled locations are expected to transition to different Köppen-Geiger climate classes by 2050, with distinct shifts observed among North American and Iranian populations. Genome environment association (GEA) analysis identified replicated temperature and precipitation associated loci across multiple chromosomes. Using genotype-phenotype data, genomic selection for cannabinoid traits revealed that only CBD achieved prediction accuracies exceeding 0.5, consistent with a polygenic architecture extending beyond the CBDAS locus. Collectively, these results demonstrate that feral and landrace Cannabis populations harbor substantial adaptive variation and represent an underutilized reservoir for climate resilient breeding and allele discovery, with relevance for pre-breeding efforts under future climate scenarios.

## Introduction

### Species Biology and Domestication of Cannabis

Cannabis (*Cannabis sativa* L.) is a wind pollinated annual species characterized by high levels of standing genetic variation and a long history of human mediated dispersal (Clarke & Merlin., 2013, Ren et al., 2021, Balant et al., 2025, Aina et al., 2025). The species is primarily dioecious although monoecious forms have been repeatedly selected in fiber hemp breeding programs, facilitating both outcrossing and rapid population expansion (Baek & Vergara., 2025). High fecundity, effective pollen dispersal (Nimmala et al., 2024) and seed dormancy contribute to large effective population sizes and promote gene flow across landscapes (Ellstrand et al., 2013) and are features that distinguish Cannabis from many other crop species. Domestication of Cannabis has resulted in a continuum of forms, selected for fiber, seed or resin production (Long et al., 2017, Rull., 2022, Del Martello et al., 2024). Selection for fiber traits favored tall, unbranched plants with long internodes and synchronous flowering, while selection for resin production emphasized compact architecture, increased branching and enhanced cannabinoid biosynthesis (Cherney & Small, 2016, Clarke & Merlin., 2013). These divergent selection pressures have produced recognizable hemp and drug-type lineages, however extensive hybridization and shared ancestry often affects strict taxonomic boundaries (Sawler et al., 2015, Lynch et al., 2016, Schwabe & McGlaughlin., 2019, McPartland & Small., 2020, Halpin-McCormick et al., 2024, Lynch et al., 2025). Cannabis is prone to ferality, with cessation of cultivation leading to escaped populations which persist in disturbed habitats (Small 1975, McPartland and Small., 2020, Aina et al., 2025). Over decades, these feral populations experience natural selection under local climatic conditions while remaining partially connected through gene flow (Wu et al., 2024). This combination of local adaptation and admixture makes Cannabis well suited to landscape genomic approaches that seek to disentangle adaptive variation and demographic history. As Cannabis differs from many annual crops, in that feral populations are dynamic and evolving, they provide a unique opportunity to study adaptation across broad climatic gradients and to identify alleles relevant for crop improvement under changing environmental conditions.

### Cannabis History in North America

Cannabis was introduced to North America early in the 17th century (Fike et al., 2016). Adoption across North America during the colonial era was widespread, being an important export and local commodity for fiber-based products (Wilsie., 2014). Early U.S. hemp introductions and cultivation practices were documented by USDA scientists such as Lyster H. Dewey in the early 20th century during a period when the United States was a major hemp producer (Dewey., 1906). This use led to increased acreage during the 19th and early 20th centuries in the United States with a peak of 176,000 acres being grown in 1943 during World War II (Wilsie and Black., 1944). Production was centered in the Midwest, bolstered by early 20^th^ century breeding efforts for yield and quality (Robinson, 1996, Schoenrock., 1966). However, despite the extent of the acreage grown, the 1937 Marihuana Tax Act eroded the value making the crop unprofitable to produce and leading to a steady decline in acreage. Much of the early agronomic and breeding knowledge was lost after prohibition, necessitating modern rebuilding of the science (Smart., 2021). This extended period of cultivation, coupled with prolific seed shatter and long distance transportation of plant material, led to many populations escaping in regions of major production prior to prohibition (Haney & Kutscheid., 1975).

### Feral Cannabis Breeding and Conservation Potential

Feral hemp populations represent a largely untapped genetic resource with significant potential to reduce germplasm vulnerability and strengthen contemporary hemp breeding programs. These populations persist across diverse landscapes, including abandoned agricultural fields, transportation corridors and farmsteads, where they have survived decades of selection under local biotic and abiotic pressures following the cessation of formal cultivation during prohibition. As a result, feral hemp likely harbors adaptive alleles related to stress tolerance, pest and disease resistance, photoperiod response, and environmental resilience that are poorly represented in modern commercial cultivars (Woods et al., 2023). Strategic collection efforts that prioritize expansion into new ecoregions and target genetically heterogeneous populations can substantially broaden the U.S. hemp gene pool. Characterizing existing collections of feral hemp will not only safeguard domestically adapted diversity but also facilitate international germplasm exchange, positioning the U.S. as an active contributor to global hemp improvement.

To accelerate breeding, access to extensive and diverse germplasm is foundational for achieving sustained genetic gain and mitigating risks associated with narrow genetic bases, a principle long recognized in crop improvement (Harlan and de Wet 1971; Byrne et al., 2018). Hemp growers currently face challenges including pest and disease pressure, cannabinoid non-compliance, poor uniformity, and inadequate regional adaptation, many of which can be addressed through breeding strategies that leverage broader genetic variability (Sunoj Valiaparambil et al., 2023). Incorporating feral germplasm into pre-breeding pipelines can introduce novel variation while reducing reliance on a limited set of international or proprietary sources (Dwivedi et al., 2023). In this context, feral hemp functions analogously to crop wild relatives in other systems (Warschefsky et al., 2014), offering resilience traits shaped by long-term natural selection. Systematic conservation of these populations through public gene banks, particularly the USDA-ARS National Plant Germplasm System, is therefore critical to reducing germplasm vulnerability.

### Landscape Genomics

There is extensive evidence demonstrating the influence of environmental factors on trait expression in many crop systems (Chapin et al., 1993, Harrigan et al., 2009, Munkvold et al., 2013). Feral cannabis populations have therefore likely had an opportunity to adapt to local environments over each generation since their escape. These populations have spread widely across North America and are established in a range of diverse climates (Wenger et al., 2020, Busta et al., 2022, Ford et al., 2024, Aina et al., 2025). This decades-long time period has allowed natural selection to act, creating an opportunity to apply genome-environment association (GEA) approaches to investigate and explore adaptation to climatic and edaphic conditions within germplasm (Campbell et al., 2025). Agricultural landscape genomics across a range of crops has shown that promising genetic variation exists (Campbell et al., 2025, Halpin-McCormick et al., 2025). Eurasian feral and landrace Cannabis have shown to be adapted to unique environments (Halpin-McCormick et al., 2025), however, a knowledge gap exists around the potential of North American material to provide additional climate adaptation.

Numerous genotype-environment association (GEA) methods have been developed to characterise genetic variation associated with environmental gradients and to infer adaptive responses to climate change (Coop et al., 2010, Forester et al., 2018, Rellstab et al., 2015). These approaches differ in their expectations and in the types of biological insights they provide. For example, some methods focus on identifying genomic regions associated with environmental variables (e.g. WZA - Booker et al., 2024), others estimate adaptive value of accessions (e.g. EGS - Campbell et al., 2025, Halpin-McCormick et al., 2025), while others are designed to guide climate matching and assisted movement of populations (e.g. FIGS - Street et al., 2016). Integrating these perspectives allows adaptation to be examined across genomic, individual and geographic scales, providing a more comprehensive basis for breeding, conservation and climate resilient germplasm deployment.

To identify *Cannabis* populations and genotypes with potential advantages under abiotic stress we combined georeferenced North American and Eurasian samples to increase statistical power, quantified variation in adaptive potential using environmental genomic estimated adaptive values (GEAVs) and assessed signatures of local adaptation using genome-environment association (GEA) approaches. These results were further contextualised using Köppen-Geiger climate classifications (Beck et al., 2018) to evaluate how current and future climate niches may change in the future. Together these analyses provide a framework for identifying candidate breeding lines and populations that may warrant conservation under projected climate change.

## Materials and Methods

### Plant Material and Genotyping

Previously, 760 feral cannabis samples were collected across twelve U.S. states and sequenced with genotyping-by-sequencing (GBS) using the restriction enzyme ApeKI (Aina et al., 2025). Raw fastq sequences were trimmed to remove Illumina TruSeq adaptor sequences using cutadapt v4.2 (1_fastq_adaptor_trim.slurm). Samples were demultiplexed using Stacks (v2.62) (2_fastq_demultip.slurm). We used the CBDRx (cs10 v.2.0) (GCF_900626175.2_cs10_genomic.fna) as the reference genome (Grassa et al., 2021). Reads were mapped using bwa (v0.7.18) (Li and Durbin., 2010). The resulting SAM file was converted to BAM format using samtools (v1.12) (Danecek et al. 2021) with a quality threshold of 10. Duplicate reads were removed from the BAM file using MarkDuplicates in Picard (http://broadinstitute.github.io/picard). Labeling of read groups was corrected using AddOrReplaceReadGroups in Picard (v3.1.0). The reference genome was indexed using samtools faidx and a sequence dictionary was created for integration with the Genome Analysis Toolkit (GATK) (DePristo et al., 2011). Using RealignerTargetCreator and IndelRealigner in GATK (v3.8) (DePristo et al., 2011) local realignment around insertions and/or deletions (indels) was performed. Using bcftools (v1.11) (Danecek et al., 2021) BAM files were merged and variant calling was performed. For filtering, variants were filtered using bcftools filter with a mapping quality of 30 (MQ < 30). Additional filtering steps were performed using VCFtools (v0.1.16) (Danecek et al., 2011) and included the following criteria (i) restriction to biallelic variants (--max-alleles 2), (ii) setting a minimum read depth (DP) between 4 and 50 (--minDP 4, --maxDP 50), (iii) filtering out genotypes with a genotype quality (GQ) below 10 (--minGQ 10), (iv) excluding variants with a quality score (QUAL) below 30 (--minQ 30), (v) removing variants with a minor allele frequency (MAF) below 0.05 (--maf 0.05) and (vi) retaining variants only if present in at least 70% of samples (--max-missing 0.7). This resulted in a total of 37,346 SNPs (Cannabis_sativa_Aina_n760_filtered.vcf.gz) (pre-filtering was 6,899,896) (Cannabis_sativa_Aina_n760.vcf.gz).

In a prior study, vcf files for the Ren et al., (2021) and Soorni et al., (2017) Cannabis datasets were generated using the same variant calling, filtering criteria and reference genome as described above (Halpin-McCormick et al., 2025). This processing resulted in 56,281 SNPs after quality filtering (Halpin-McCormick et al., 2025). When the Aina et al., (2025) feral dataset was overlapped with Ren et al., 2021 and Soorni et al., 2017 datasets there was a total of 13,389 SNPs remaining and 909 samples in the combined datasets (**Table S1**). The 13,389 SNPs were distributed across all chromosomes with a mean distance of 63,626 bp between consecutive SNPs.

### Maximum Likelihood Phylogeny

A maximum likelihood phylogeny was constructed on the 909 samples rooted using the *Humulus lupulus* reference genome (GCA_963169125.1_drHumLupu1.1_genomic.fna) (Natsume et al., 2015). The 13,388 SNP positions were mapped to corresponding positions in the *Humulus lupulus* genome using bedtools (v2.31.1) (Quinlan and Hall 2010) and the corresponding SNPs extracted. The vcf file was converted to phylip format using the vcf2phylip software (Ortiz, 2019) handling heterozygous ambiguities using the - m 1 mode and IUPAC ambiguity codes. A phylogenetic tree was inferred using IQ-TREE (v2.3.6) (Nguyen et al., 2015) with the best substitution model selected using -m MFP flag (GTR+F+ASC+R10) with ascertainment bias +ASC and -bb flag for bootstrap support. Trees were visualized in FigTree (v1.4.4) (http://tree.bio.ed.ac.uk/software/figtree/).

### Köppen-Geiger Climate Shifts

Present day (1991-2020) and future (2041-2070) Köppen-Geiger maps at 1km resolution (Beck et al., 2023) (v2) were downloaded from www.gloh2o.org/koppen. To assess how climate classes may change through time, the climate classes were extracted and counted for the 191 samples with latitude and longitude positions for both present and future periods, respectively (**Table S2**).

### fastSTRUCTURE analysis

To examine structure and shared admixture proportions, VCFtools was used to generate map and ped files for the input VCF file. PLINK (v1.90b6.21) (Purcell et al., 2007) was then used to convert these files into bed, bim and fam files. Population structure was assessed using fastSTRUCTURE (v1.0) (Raj et al., 2014). Optimal K was examined using the silhouette and elbow methods in the FactoMineR (v2.11) (Lê et al., 2008) and factoextra (v1.0.7) (Kassambara and Mundt 2016) packages.

### Climate Data

Accession level latitude and longitude coordinates enabled the extraction of location specific climate data. Three distinct datasets were included in this analysis. The Ren et al., 2021 dataset comprising 82 unique latitude and longitude points, the Soorni et al., 2017 contained 67 unique locations and the Aina et al., 2025 dataset comprising 85 unique latitude and longitude points. All climate data was retrieved from the WorldClim database (Fick and Hijmans, 2017). Climate data for these 191 unique data points can be seen in **Table S2**. Additionally, each site was classified using the Köppen-Geiger climate class system (Beck et al., 2023). Köppen-Geiger climate classes were summarized for each U.S. state across four historical (1901-1930, 1931-1960, 1961-1990, 1991-2020) and two future time points 2041-2070, 2071–2099, SSP585) (Beck et al., 2023). For each state, proportional climate class composition and the fraction of area undergoing class transitions between consecutive periods were calculated from rasters using spatial overlays, where each grid cell represents approximately a 10 × 10 km area and is assigned a single climate class (**Figure S18**). Analyses were performed in R using the terra (v1.8.86) package (Hijmans, R.J., 2025).

### Environmental Genomic Selection

#### Fusion of Ren, Soorni and Aina datasets

For the multi-dataset EGS training set, a greedy algorithm was used previously on the Aina dataset to iteratively select individuals with the greatest pairwise genetic distance. Where multiple samples occurred at the same location, a representative genotype was chosen at random. This resulted in coverage of 80/85 potential population lat/long points. The full training set was therefore 80 samples from the Aina dataset, 67 samples from Soorni dataset and 44 samples from Ren dataset. Therefore 191 samples (∼21 %) with latitude longitude positions were used as the training set for the multi-dataset EGS (**Table S2**), leaving 718 samples in the test set. Overlapping the three datasets as stated above resulted in 13,389 common SNP sites. Prediction accuracies for the 19 WorldClim bioclimatic traits were calculated similarly to the above for the combined datasets (**Table S3**) and were calculated on 12,030 SNPs as this was the SNP count returned across the 191 training samples when filtered for SNPs less than or equal to 20 % missingness. Filtering to remove SNPs with greater than 20 % missingness across all samples left a final SNP set of 12,959. Environmental genomic selection (EGS) with the rrBLUP model was performed using 12,959 SNPs, with 7.9 % missingness imputed with the marker mean (5_Genomic_Selection.R) (**Table S4**).

#### Aina dataset

A training set of 310 (∼40 % of the dataset) was developed by maximizing genetic distance for sample selection across all 760 samples (1_core_identification.py) (**Table S5**). Five samples within the 310 sample training set had no environmental data and were removed, with a final training set of 305 (**Table S5**) and 455 samples in the test set. The vcf (Cannabis_sativa_Aina_n760.vcf.gz) was converted to rrBLUP format (vcf_to_rrBLUP_format.py). Further filtering to remove SNPs with greater than 20 % missingness across all samples left a final SNP set of 22,852. This filtering step left a genome wide average missingness per sample of 13.3 %, this was then imputed with the marker mean. Prediction accuracies for the 19 WorldClim bioclimatic traits was calculated (4_Cross_validation.R) (**Table S6**) with four genomic prediction methods; RR-BLUP, G-BLUP with an exponential kernel, G-BLUP with a Gaussian kernel, and BayesCπ. The R packages ‘rrBLUP’ (Endelman, 2011) and ‘hibayes’ (Yin et al., 2022) were used for the analysis. Prediction accuracies were calculated on 26,433 SNPs as this was the SNP count returned in the 305 training samples when filtered for SNPs less than or equal to 20 % missingness. This filtering step left a genome wide average missingness per sample of 8.3 % that was imputed with the marker mean. Prediction accuracy was based on Pearson correlation (r(PGE,y)) between the predicted genotypic effects and the observed environmental variable with 10-fold cross-validation. Environmental genomic selection (EGS) with the rrBLUP model was performed using 22,852 SNPs (5_Genomic_Selection.R) (**Table S7**).

### Genomic Selection

Within the Aina dataset, secondary metabolite data was collected for cannabinoid traits namely; CBG, CBD and Δ9-THC and total cannabinoid content (sum of eight cannabinoids; CBC, CBD, CBDV, CBG, CBN, Δ8-THC, Δ9-THC, THCV). A total of 276 samples with complete chemistry data (**Table S8**) were used as the training set while the remaining 484 samples were used as the test set. GS using 37,346 SNPs was performed with 14.65 % missingness imputed by the marker mean at these NA sites. Prediction accuracy using the rrBLUP model was calculated for the cannabinoid traits (**Figure S14A, Table S9**). GEBVs were calculated for four cannabinoid traits (**Figure S14B, Table S10**). The genomic distribution of large-effect markers (top 1 % by absolute rrBLUP marker effect size; n = 374) was examined. Top 1 % SNPs were annotated based on overlap with transcript features where possible. SNPs falling outside annotated transcripts were assigned to the nearest transcript to facilitate functional interpretation (**Table S11**).

### Genome Environment Association Analysis

To test for genome-environment association, genotypes were collapsed into population level allele counts. The drug-type population was removed due to lack of environmental data leaving ten subpopulations and 13,389 SNPs and 19 WorldClim environmental variables. Allele counts per SNP and population were calculated. Kendall’s τ rank correlation was performed between allele frequency and each environmental covariate across populations, yielding per SNP p-values. Weighted Z-analysis (WZA) was applied combining SNP level p-values into window (50k per chromosome) scores (Booker, 2024). SNPs surpassing the WZA significance threshold were extracted and annotated, with results reported separately for temperature-related (**Table S12**) and precipitation-related (**Table S14**) bioclimatic variables.

### Spatial ancestry inference using TESS3

Spatial ancestry was inferred using the tess3r R package (v1.1.0) (Caye & François., 2016) with geographic coordinates incorporated into the model and ancestry coefficients visualised using kriging based spatial interpolation across North America and Eurasia.

### Growth Chamber Experiment

#### Plant Material and Experimental Design

Seven hemp (*Cannabis sativa* L.) accessions (**Table S16**) were evaluated for early growth responses under controlled greenhouse and growth chamber environments. Seeds of each accession were sown into 3 × 3 inch plastic pots filled with PRO-MIX potting soil. A total of 12 replicates per accession were established. All plants were grown in the Walnut Street Greenhouse at the University of Wisconsin-Madison using a randomized complete block design. Greenhouse conditions were maintained at approximately 25 °C with a 14 h light and 10 h dark photoperiod. Plants received daily overhead irrigation. At seven days after planting, plant height was measured in centimeters from the soil surface to the apical meristem for all replicates.

#### Heat Treatment and Growth Chamber Conditions

Following the initial phenotyping, six randomly selected replicates per accession were transferred to a GEN1000 growth chamber (Conviron, Manitoba, Canada) for heat stress treatment. Plants were arranged in a randomized block design within the chamber. Environmental conditions were set to 35 °C, a 14 h light and 10 h dark photoperiod, ∼100 μmol m^−2^ s^−1^ light intensity, and 40 % relative humidity. The remaining six replicates for each genotype remained in the greenhouse under the original conditions.

#### Phenotyping

After seven and 14 days of exposure to their respective environments, plant height was recorded again in centimeters for all surviving plants.

## Results

### Data Merging and Population Structure

Three datasets, Ren et al., 2021 (n = 82), Soorni et al., 2017 (n = 67) and Aina et al., 2025 (n = 760) were merged as they contained genotyped and georeferenced individuals (**Figure 1A, Table S1**). Raw sequence data from all studies were used in a common pipeline to call SNPs resulting in a combined dataset of n = 13,389 SNPs across the genome. Among the merged population (n = 909) there were 191 unique georeferences and substantial population structure that was largely geographic, with 11 populations observed (**Figure 1**) and broadly corresponding to collection sites (**Figure 1B/C**).

**Figure 1.**
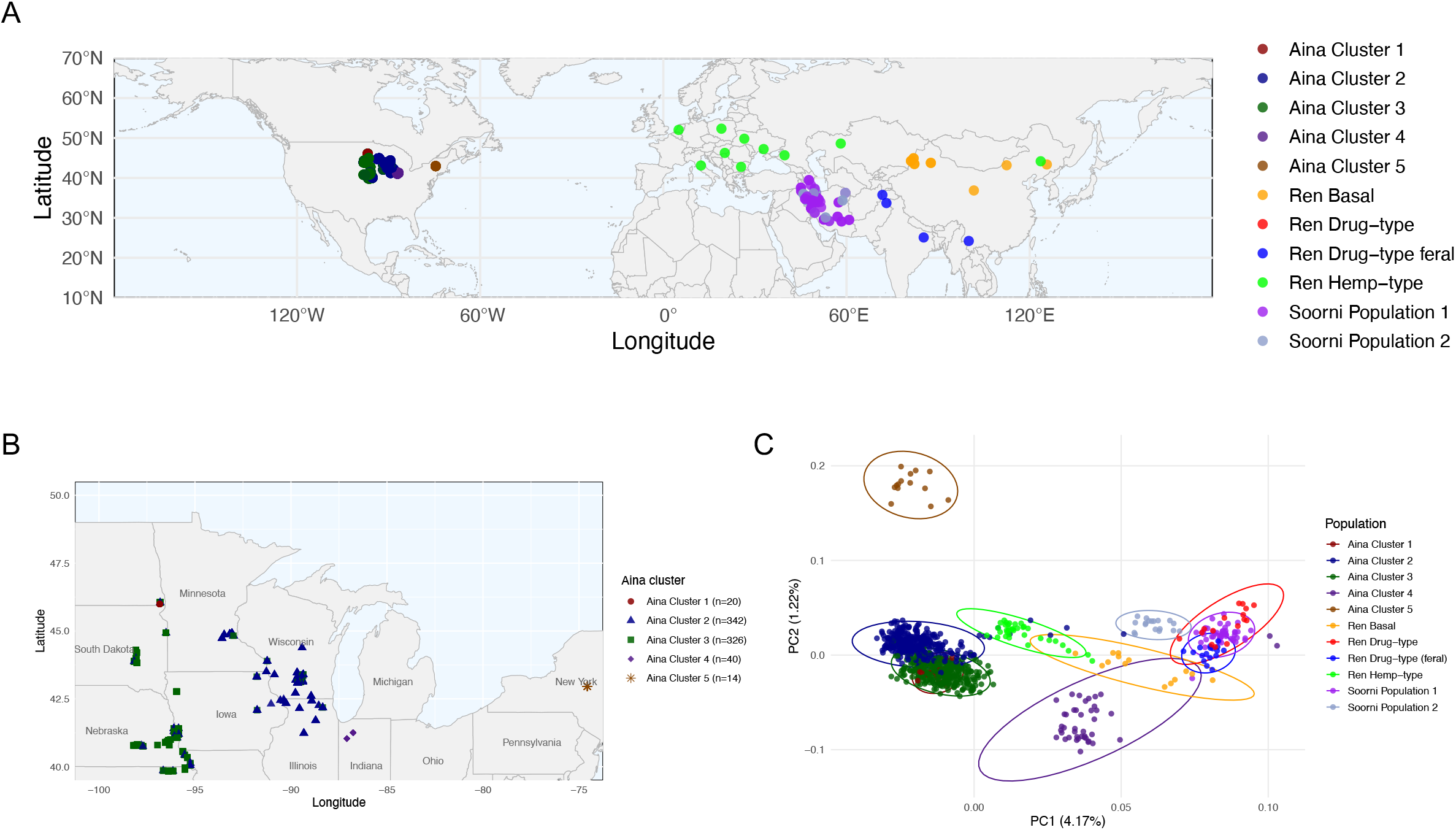
Fusion of Eurasian and North American Cannabis datasets revealed 13,389 overlapping SNPs (**A**) Global population distribution of samples with 191 unique latitude and longitude points (**B**) Population distribution of North American (n=760) samples (**C**) PCA on 909 samples (Ren = 82, Soorni =67 and Aina = 760) with 2,792 SNPs following LD pruning (LD=0.2).

### Relationship between North American and Eurasian Collections

As the relationship between the North American and Eurasian collections was unknown, a maximum likelihood phylogenetic tree (n = 13,388 SNPs) rooted to *Humulus lupulus* was generated. Phylogenetic structure broadly reflects group relationships associated with geographic collection sites (**Figure 2**). North American accessions (Aina cluster 1-5) showed a closer and well supported genetic relationship to basal (orange) and hemp-type (bright green) lineages (bootstrap = 98) rather than to drug-type feral (blue), drug-type (red) or Iranian populations (purple and teal). U.S. Drug-type accessions (red) were observed to cluster more closely with drug-type feral (blue) and Iranian populations (**Figure 2**). This was additionally supported in fastSTRUCTURE analysis (**Figure S1**) with the same 13,389 SNP set, where K=2 separated the North American samplings (Aina Clusters 1-5) and European hemp-type from drug-type, drug-type feral and Iranian Population 1 samples, with Iranian Population 2 and basal samples intermixed (**Figure S1C**). At K=4-6, Aina Cluster 4 (Indiana, dark purple) and Aina Cluster 5 (New York, brown) samples show relatively low admixture and increased assignment consistency, suggesting greater genetic distinctiveness and consistent with their increased separation observed in PCA space (**Figure 1C**).

**Figure 2.**
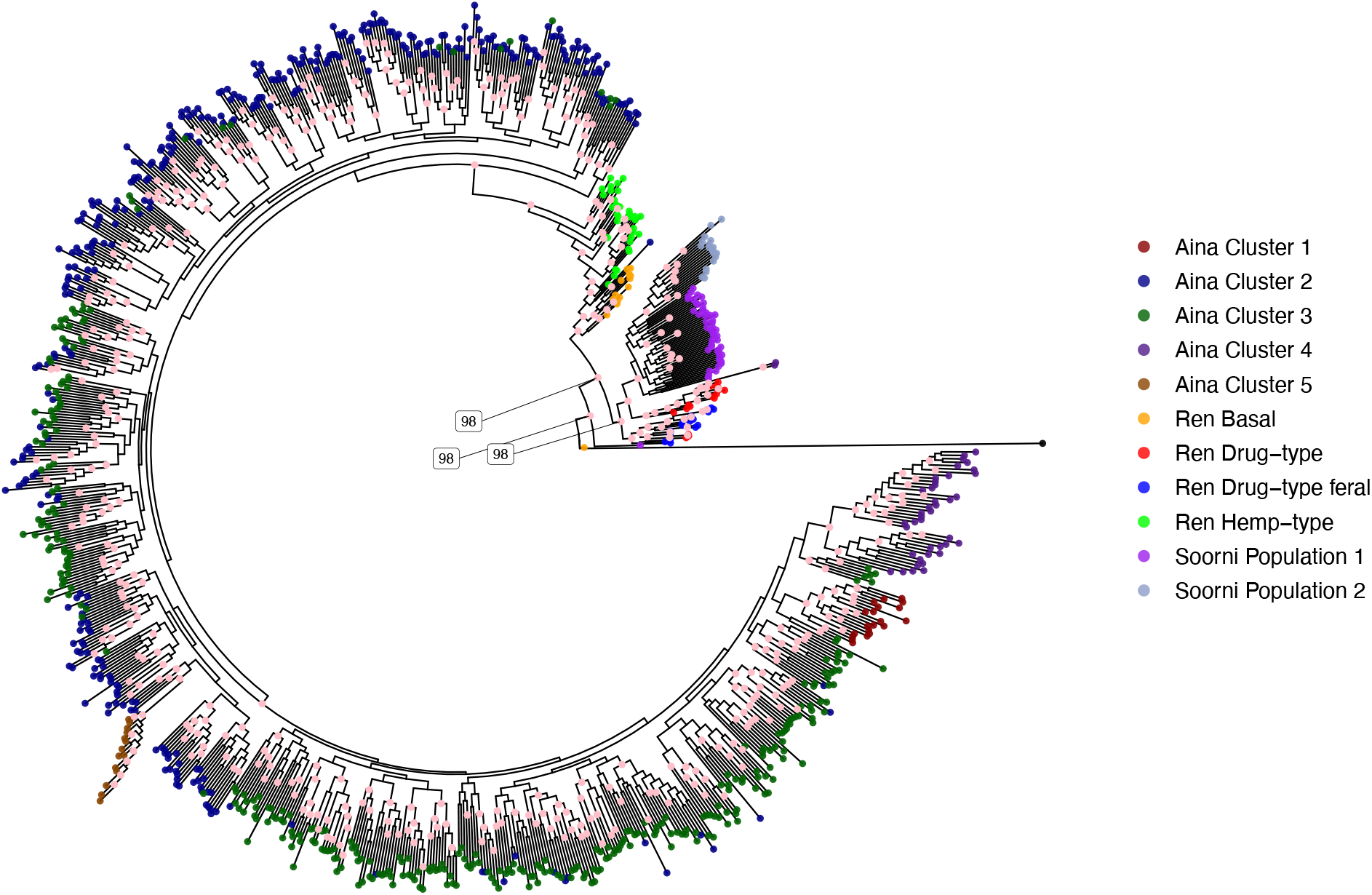
Maximum likelihood tree for 909 samples and 13,388 SNPs (1 invariant sites removed) rooted to *Humulus lupulus*. Using IQTree, ModelFinder selected GTR+F+ASC+R10 as the best-fit model based on AIC, corrected AIC and BIC. This model assumed General Time Reversible (GTM) substitution with ascertainment bias (ASC) due to SNP only data with rate heterogeneity across sites using FreeRate model with 10 rate categories (R10). Ultrafast boot strap analysis with 1,000 replicates was conducted to assess node support. Local rearrangements using Nearest Neighbor Interchange (NNI) were performed on each bootstrap replicate. Of 908 internal nodes 451 (49.7%) received greater than 95% bootstrap support, indicating moderate overall confidence in tree topology. The 451 nodes with >95% support are indicated in pink. The first inner nodes bootstraps of 98 are indicated. To aid visualization, the terminal branch of the *Humulus lupulus* outgroup was shortened (scaled to 20% of its original length), this adjustment does not affect tree topology or relative branch lengths among Cannabis samples.

Spatial population structure was further supported by tess3r, which interpolates ancestry coefficients across geographic space (**Figure S2**). At K=5 and K=6, tess3r revealed clear spatial differentiation among the North American samples (Aina Cluster 4 (Indiana) & Aina Cluster 5 (New York)) and consistent separation of European and Chinese samples from Iranian collections in Eurasia (**Figure S2B**). Genome wide nucleotide diversity varied among populations, with North American Aina Cluster 5 (New York) exhibiting a consistently lower diversity and increased chromosome specific variation relative to other population groups (**Figure S3**).

### Environmental Genomic Prediction

Environmental genomic prediction was conducted on the combined North American and Eurasian collections. A training population of 191 samples corresponding to unique latitude and longitude coordinates was used across 19 bioclimatic variables (**Table S2**), with predictions generated for the remaining 718 samples in the dataset. Prediction accuracy was assessed using 10-fold cross-validation with 50 iterations across four models (**Figure S4, Table S3**). Nine bioclimatic variables exceeded the 50 % accuracy threshold in prediction accuracy (**Figure 3A, Figure S4 & Table S3**). Environmental genomic selection (EGS) results on the combined Eurasian and North American Cannabis datasets (n=909) showed marked differences in genomic estimated adaptive values (GEAVs) across the 11 population groupings (**Figure 3A, Figure S5 & Table S4**). Overall, North American populations tended to exhibit positive GEAVs for precipitation related variables, such as seasonal and annual precipitation (eg. Bio 8 and Bio 12) (**Figure 3A**), consistent with adaptation to wetter environments. In particular, Aina Cluster 5 (New York) showed elevated GEAVs for precipitation during the driest and coldest quarters (Bio 17 and Bio 19) and precipitation during the driest month (Bio 14), while exhibiting a comparatively lower GEAV for maximum temperature of the warmest month (Bio 5) (**Figure 3A**), highlighting contrasting adaptive signatures across climatic gradients experienced by populations.

**Figure 3.**
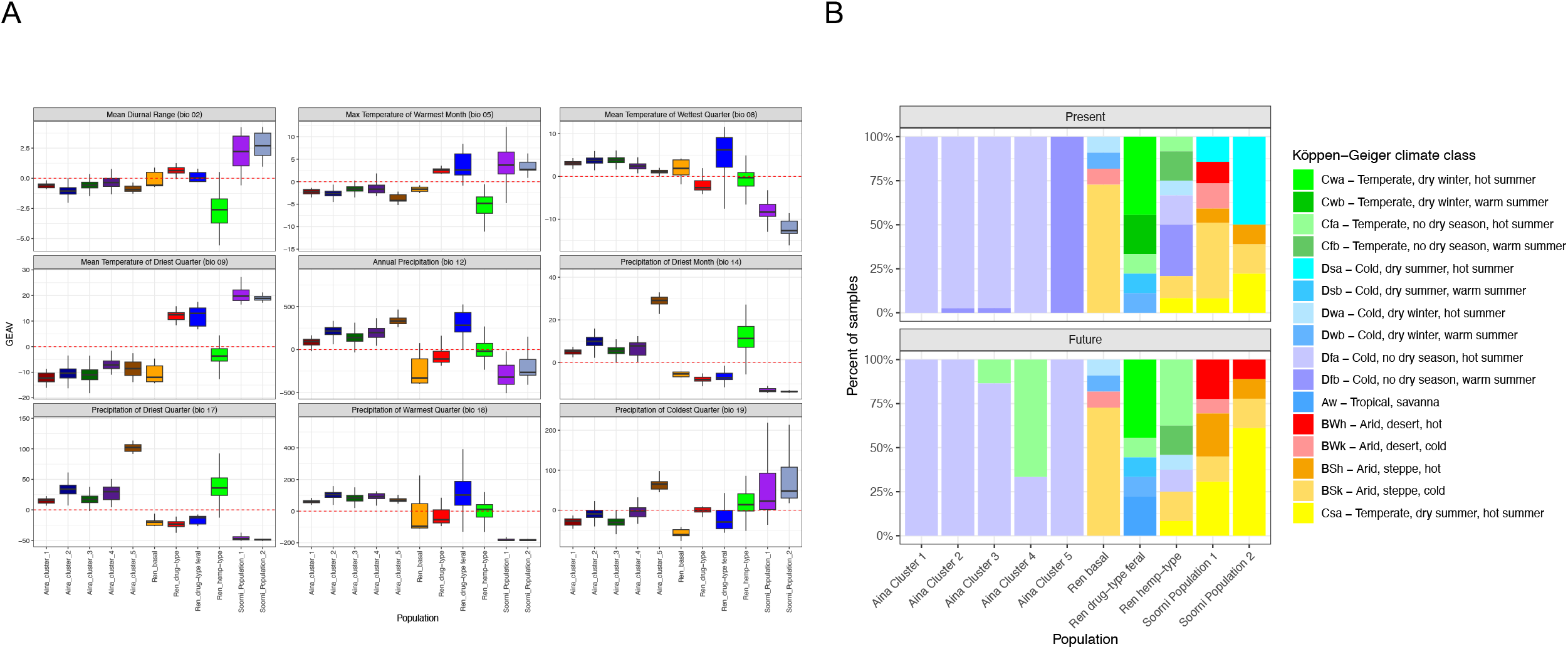
EGS on Eurasian and North American Cannabis datasets (n=909) for the 11 population groupings (**A**) GEAV results for the 9 WorldClim bioclimatic traits with prediction accuracies above 0.5 (Bio 2, Bio 5, Bio 8, Bio 9, Bio 12, Bio 14, Bio 17, Bio 18 and Bio 19) (**B**) Changes in Köppen-Geiger climate classes for the 191 samples with latitude and longitude values, grouped by population, for the present day (1991–2000) and the future period (2041–2070) under SSP585.

The 191 samples with unique latitude-longitude coordinates spanned 15 Köppen-Gieger climate classes (**Figure 3B, Figure S6 & Table S2**). Looking from the present day (1991-2000) to the future (2041-2070) under the SSP585 or high emission climate change scenario, we see climate class representation losses in the more under-represented climate classes (12 - Cwb - Temperate, dry winter, warm summer, 17 - Dsa - Cold, dry summer, hot summer; 26 - Dfb - Cold, no dry season, warm summer) while temperate type climate classes show increases (8 - Csa - Temperate, dry summer, hot summer, 14 - Cfa - Temperate, no dry season, hot summer), with 126 samples (∼ 66 %) showing no change and 65 samples (∼34 %) showing a change in climate class at that location moving into the future (**Figure S6B, Table S2**).

Future climate projects also indicate population specific responses. Notably, Aina Cluster 4 (Indiana) is projected to transition away from historically cold, no dry season and hot summers (Dfa) to warm temperate regimes (Cfa), while Aina Cluster 5 (New York) shows a change from cold, no dry seasons, warm summers (Dfb) to cold, no dry season, hot summers (Dfa) (**Figure 3B**). The two populations from the Soorni Iranian collection show the strongest shifts overall, with clear increases in Arid and dry summers (BWh) and temperate climate class (Csa) and losses in cold, dry summer, hot summers (Dsa). Responses were also evident for the Ren dataset, with drug-type feral populations showing losses in temperate, dry winter, warm summers (Cwb), whereas hemp-type show increases in temperate, dry winter, warm summer (Cfa) climate classes in the future projection (**Figure 3B**).

The Aina et al., 2025 dataset alone included 760 samples from 85 unique locations (**Table S5**) and consisted of 37,346 SNPs post filtering. A principal component analysis following LD pruning (0.2) resulted in 3,762 SNPs and reproduced population structure consistent with previous publication (Aina et al., 2025) (**Figure S7A**) with five populations namely; Cluster 1 (West North Central-b, WNCb, dark red), Cluster 2 (Mississippi River, dark blue), Cluster 3 (West North Central-a, WNCa, dark green), Cluster 4 (Indiana, dark purple) and Cluster 5 (New York, brown). fastSTRUCTURE analysis using 37,346 SNPs further supported these results, with K=5 identified as the optimal cluster number based on the elbow and silhouette methods (**Figure S8A**). fastSTRUCTURE analysis at K=5 revealed clear separation of Clusters 2 - 5 into distinct ancestry components, while Cluster 1 (WNCb, dark red) remained admixed and was not resolved as an independent cluster (**Figure S8C**). This is additionally observed in the principal component analysis where Cluster 1 (WNCb, dark red) is centrally positioned and overlaps with Cluster 2 (Mississippi river, dark blue) and Cluster 3 (WNCa, dark green) (**Figure S7A**). At K=5 Cluster 4 (Indiana, dark purple) and Cluster 5 (New York, brown) form well defined ancestry groups with limited admixture and clear separation from the remaining clusters, similar to that seen in the combined dataset (**Figure S1C, Figure 1C**).

Using a de-novo core collection that maximized genetic distance among individuals as a training population (n = 305) (**Figure S9**) we similarly conducted an environmental genomic prediction on 19 bioclim variables for the Aina et al., 2025 dataset alone. Prediction accuracy was assessed similarly, with 10-fold cross-validation and 50 iterations across four models, using 26,433 SNPs (**Figure S10, Table S6**). When looking only at the Aina dataset alone, prediction accuracies were high, likely due to the high number of individuals that were sampled from the same location (**Figure S10**). GEAVs for the Aina dataset were estimated across the 19 bioclimatic variables using 22,852 SNPs (**Figure S11, Table S7**), with fourteen variables exceeding the 50 % accuracy threshold in prediction accuracy (**Figure S12, Table S6**). GEAV pattern for Bio 14 and Bio 17 for Cluster 5 (New York) were consistent across datasets, with similar trends observed across populations between the Aina alone dataset using 22,852 SNPs and the reduced 12,959 SNPs used for the GEAV estimated from the combined datasets (**Figure S12 & Figure 3A**).

### Genomic Prediction for Cannabinoid Traits

The availability of paired genotype and cannabinoid phenotype data within the Aina dataset enabled the application of a conventional genomic selection (GS) approach to calculate prediction accuracies (**Figure S13A, Table S9**) and genomic estimated breeding values (GEBVs) for cannabinoid traits (**Figure S13B, Table S10**). GEBVs for four cannabinoid traits were estimated using 37,346 SNPs, with 273 samples in the training set and 487 samples in the test set (**Figure S13, Table S10**). Among the four traits evaluated, only CBD achieved a prediction accuracy exceeding the 0.5 threshold (**Figure S13A**). Despite variable prediction accuracy across traits, population level differences in GEBVs were evident for total cannabinoids, CBG, CBD and Δ9-THC (**Figure S13B**). Results suggest that while major loci are known to contribute substantially to CBD variation (de Meijer et al., 2003, Weiblen et al., 2015, Laverty et al., 2019, Wenger et al., 2020, Grassa et al., 2021), additional small-effect loci distributed across the genome may also contribute to CBD potency and can be captured by the GS model (**Figure 13**).

To assess the genomic basis of CBD genomic estimated breeding values, the top 1% of SNPs by absolute marker effect size (n=374), including both positive and negative effects were examined (**Figure S14**). Marker effects were distributed non-uniformly across the genome, with nine SNPs clustering within the CBDAS region on chromosome 7 (29.6-61.4Mb) and revealing putative effect hotspots (end of chromosome 1, start of chromosome 10) (**Figure S14**). Full annotations for these top ranked markers are provided in **Table S11**. The annotated transcripts were enriched for genes involved in stress and defense signalling, hormone transport and signalling and metabolic regulation, rather than direct cannabinoid biosynthetic enzymes. Notably, multiple high-effect SNPs were associated with transcripts encoding NLR-type disease resistance proteins, auxin transporters and calcium-dependent kinases, supporting a highly polygenic architecture underlying CBD content (**Table S11)**.

### Genome Environment Associations

To identify genomic regions associated with climatic variation across the species range, genome-environment association (GEA) analyses were performed using population-level allele frequencies and WorldClim bioclimatic variables, excluding drug-type samples from the Ren dataset due to their likely indoor cultivation and limited environmental relevance (**Figure 4, Figure S15 & Figure S16**). A total of 434 unique SNPs exceeded the WZA significance threshold of −log10(p) ≥ 2 (p ≤ 0.01) for temperature genome-environment associations (**Figure S15**). Annotations for these temperature associated GEA SNPs were obtained from the cs10 reference genome (**Table S12**).

**Figure 4.**
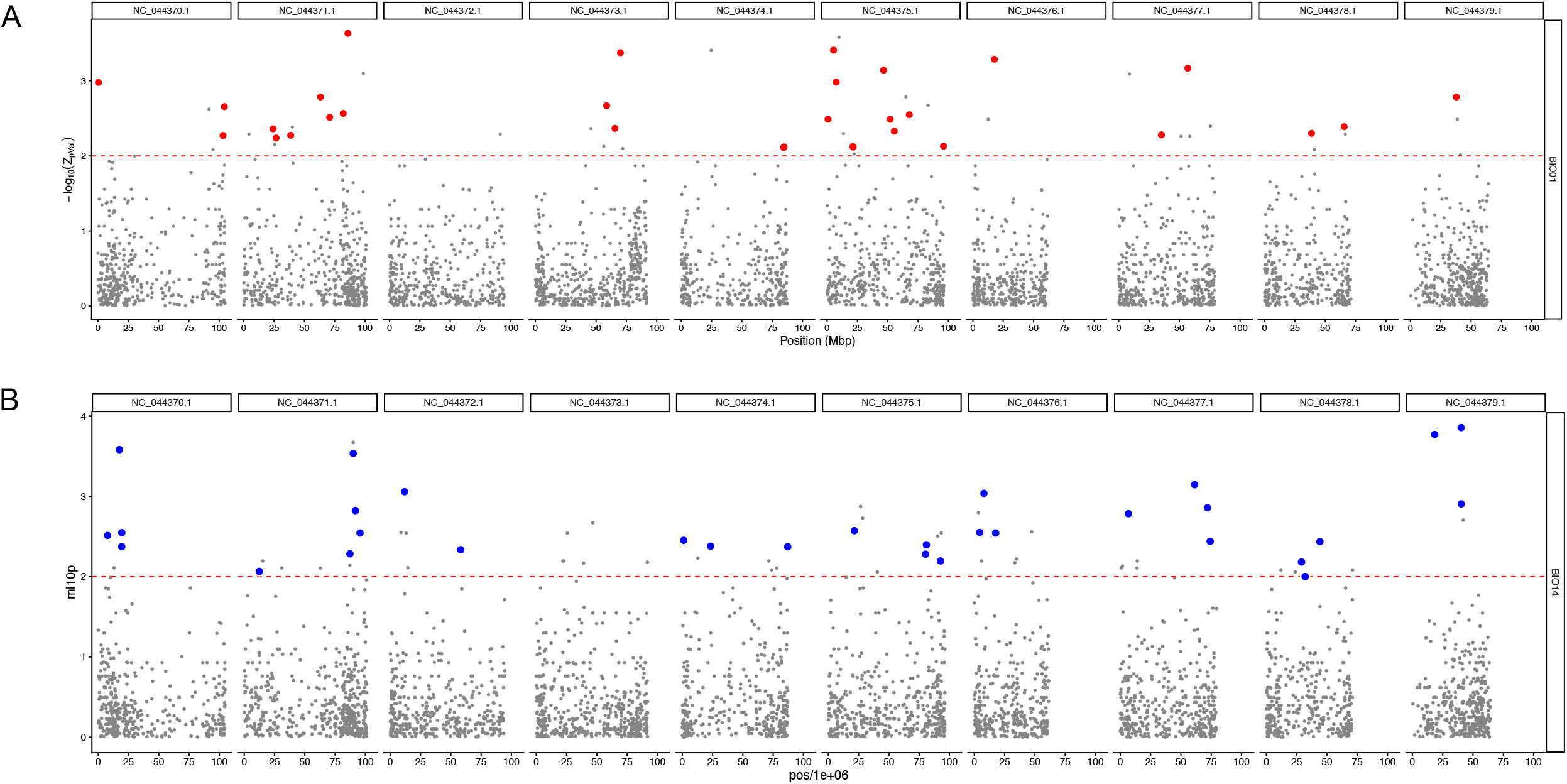
Genome-environment associations (GEA) using Kendall’s τ between population allele frequency for the WorldClim variables across ten populations. SNP-level p-values were aggregated into 50-kb window scores using the Weighted Z-analysis (WZA). The red dashed line represents -log10 (0.01) (**A**) GEA results for Bio 1 temperature (Annual Mean Temperature). Red points indicate loci that exceeded the WZA significance threshold in at least three temperature-related WorldClim variables (Bio 1 - Bio 11) (**B**) GEA results for Bio 14 (Precipitation of the Driest Month). Blue points indicate loci that exceeded the WZA significance threshold in at least three precipitation-related WorldClim variables (Bio 12 - Bio 19).

To prioritise robust temperature associated loci, SNPs exceeding the WZA threshold and observed in three or more temperature related bioclimatic variables (Bio 1 - 11) were retained (n=52, **Table S13**). Although no single climatic variable produced pronounced genome-wide peaks, several genomic regions showed consistent replication across multiple temperature-related variables (frequency ≥ 3), suggesting shared underlying temperature-associated signals. For example, loci on chromosome 2 (NC 044371.1) and chromosome 6 (NC_044375.1) repeatedly exceeded the WZA significance threshold across Bio 1, Bio 6, Bio 9, and Bio 11, indicating a recurrent association with temperature variables captured by multiple climatic descriptors (**Figure 4A, Figure S15**). Loci replicated across multiple temperature variables (Bio 1, Bio 6, Bio 9 and Bio 11) for chromosome 6 (NC 044375.1) were annotated as genes involved in transcriptional regulation (bHLH3, position 5315096), ribosome biogenesis (LSG1-2, position 7576483) and cellular trafficking (BEACH domain–containing protein B, position 46562056). The repeated detection of these loci across distinct temperature variables suggests general thermal tolerance and cellular stress response mechanisms rather than season specific climatic factors (**Figure 4A, Table S13**).

Similarly, a total of 316 unique SNPs exceeded the WZA significance threshold of −log10(p) ≥ 2 (p ≤ 0.01) for precipitation associated (Bio 12 - 19) genome-environment associations (**Figure S16**), with annotations for these examined (**Table S14**). Similar to the results for temperature, no single precipitation climatic variable produced pronounced genome-wide peaks, however several genomic regions showed consistent replication across multiple variables (frequency ≥ 3). A total of 49 SNPs exceeded the WZA threshold and were observed in three or more precipitation related bioclimatic variables (Bio 11 - 19) (**Table S15**). Replicated precipitation associated loci were most frequently detected across Bio 14 (precipitation of the driest month), Bio 15 (precipitation seasonality) and Bio 17 (precipitation of the driest quarter) (**Figure S16**). In particular, replicated SNPs exceeding the WZA threshold were observed toward the end of chromosome 2 (NC_044371.1), the end of chromosome 6 (NC_044375.1) and the start of chromosome 7 (NC_044376.1) suggesting shared precipitation-associated signals across these regions (**Figure 4B**).

## Discussion

### Limitations

This analysis is subject to several limitations, including those related to sequencing. The Aina and Soorni datasets were genotyped using GBS, which reduces the number of markers relative to the Ren dataset, which was genotyped with WGS. A second limitation is the number of latitude and longitude positions (n = 191 georeferences) relative to plant samples (n = 909), where expanded global sampling with paired genotype and environmental metadata would enhance the power and resolution of future analyses. The high prediction accuracies seen in the EGS performed on the Aina dataset alone (**Figure S10**) are likely inflated due to sample number relative to unique latitude and longitude positions, with 760 samples of which 742 fall into 85 population level coordinates. With respect to the genomic predictions performed here for four cannabinoid traits (**Figure S13**), training was only possible on low Δ9-THC (0.01-2.47 %) and moderate CBD (0.01-4.25 %) samples. While this is appreciable due to the complexities of permitting and regulations associated with researching Cannabis within academic institutions, it limits our understanding and potential to identify other small effect markers which may contribute meaningfully to secondary metabolite profiles within the plant. With crop compliance at the forefront of all hemp cultivators’ minds, being able to develop reliable breeding values for secondary metabolite profiles could help production license holders meet the less than 0.3 % THC requirement. The prediction accuracy seen for CBD concentration (> 0.5) (**Figure S13A**) suggests there may be other small effect contributors yet to be identified which may play a role in secondary metabolite expression. The inclusion of more broadly diverse secondary metabolite profiles and associated phenotyping would enable the development of GEBVs for secondary metabolite potentials and could help establish a 0.3 % threshold based framework for sample classification. The maximum likelihood phylogeny additionally assumes a one-to-one site correspondence between *Cannabis* and *Humulus* for rooting, an assumption that should be interpreted with great caution, especially given recent publications examining synteny between the two species (Carey et al., 2025).

### Ancestry of North American Feral Hemp

The integration of multiple independent datasets enabled a more comprehensive assessment of the ancestry of North American feral hemp than has previously been possible. By combining accessions spanning feral, basal, hemp-type, and drug-type lineages across broad geographic origins, this analysis provides a unified framework for evaluating the genetic relationships and historical admixture patterns within North American germplasm. Phylogenetic analysis supports shared ancestry between North American accessions with basal and hemp-type lineages (bootstrap support 98) (**Figure 2**). This clustering is consistent with historical introductions of fiber and seed hemp to North America (Cherney & Small, 2016) and suggests that a substantial proportion of feral populations retain genetic affinity with hemp gene pools. In contrast, U.S. drug-type accessions grouped more closely with drug-type feral and Iranian accessions, indicating divergent ancestry patterns within North American germplasm (**Figure 2**). Within North America, additional populations structure was evident, with accessions originating from New York and Indiana forming distinct genetic clusters across multiple analyses, including principal component analysis, fastSTRUCTURE and phylogenetic reconstruction (**Figure 1C, Figure S1C, Figure S7A, Figure S8C, Figure 3**). The consistent differentiation of these populations across independent methods suggests the potential persistence of regionally distinct feral hemp lineages within the United States.

### GEAVs and crop vulnerability

Climate is recognized as a major driver of vegetation distribution (Woodward., 1987). Further, the reduced genetic variation in cultivated crops may limit climate resilience and adaptive capacity to future environmental conditions (Zhang et al., 2017). Plant genetic resources (PGRs) encompass all the variation present in a crop species, including modern varieties, landraces and wild relatives (Salgotra & Chauhan., 2023). The collection and preservation of PGRs in their natural habitats, where adaptation and evolution is ongoing, is imperative for securing long term crop security. Without implementing conservation practices it is likely these resources will vanish (Hammer et al., 2010, Allendorf et al., 2010). More specifically for Cannabis, ongoing research and discussions have focused on if truly wild *Cannabis* still exists (Balant et al., 2025), with some evidence supporting this (Lynch et al., 2025). The amount of extant wild material remains uncertain, thus there is an urgent need to preserve locally adapted landrace/feral material. Crop improvement through the utilization of landraces or crop wild relatives could confer resilience to changing climates and increase germplasm diversity, ensuring preservation for use within active breeding programs.

Population level genomic estimates reveal geographically structured patterns of climate associated variation. Notably, accessions from New York exhibited elevated GEAVs for precipitation related variables such as Bio 14, Bio 17 and Bio 19, showing signatures consistent with adaptation to wetter and cooler environments (**Figure 3A**). Projections of future climate conditions indicate loss of representation for several Köppen-Geiger climate classes currently occupied by these populations alongside a broader shift toward more arid climate niches across populations (**Figure 3B, Figure S6B**). Together, these results highlight a potential mismatch between existing locally adapted genetic variation and future climatic conditions (**Figure S17**), reinforcing the importance of conserving landrace diversity before such adaptive signatures are lost. Poorly represented Cannabis populations are at particular risk, as many landraces are rarely conserved in ex situ genebanks, limiting their long-term preservation and accessibility (Torkmaneh & Jones, 2022). The encroachment of improved cultivars into regions traditionally occupied by landraces has been documented in countries such as Pakistan, Afghanistan, Iran, Tajikistan, Northern India and Morocco (Torkmaneh & Jones, 2022) and may lead to the erosion or loss of locally adapted genetic diversity (Khoury et al., 2021). Similar processes may occur in North America if cultivated hemp varieties are introduced or grown in close proximity to feral populations.

### GEAV Predictions and Phenotypic Exploration of Seedling Heat Tolerance

As Bio 5 (maximum temperature of the warmest month) showed moderate but robust genomic prediction accuracy (prediction accuracy = 0.55 ± 0.003), this variable was selected for further evaluation by comparison with independent field and controlled environment phenotypic data. Full rankings of population mean GEAVs for Bio 5 show that while individuals in populations have a range of values, there are clear populations with advantages (**Figure S17A**). When GEAV predictions for Bio 5 (**Figure S17A**) were compared with phenotypic outcomes from a 2024 field experiment in Arlington, Wisconsin, partial overlap was observed between the highest-ranking GEAVs and increased plant height among the same populations evaluated in Aina et al., (2025), namely population NE-22-ID-06 and population KS-22-PB-04 (**Figure S17A/B**, sample names highlighted in red). Climatic data for the Arlington site in 2024 (43.32434°N, −89.33777°W, https://aquastat.fao.org/) indicates that conditions were drier than the long-term average (763 mm versus 934 mm annual precipitation), while temperatures were not exceptionally high.

From the ten highest and ten lowest GEAVs for Bio 5 (**Figure S17A**), seven feral accessions were used to explore the impact of temperature on different georeference lineages (**Figure S17A**, red and blue arrows) (**Table S16**). In a growth chamber experiment, accessions were exposed to control (25 °C) and high temperature (35 °C) over a two week period. At 25 °C (**Figure 17C**, upper panel), genotypes exhibit relatively parallel growth patterns with consistent separation among lines. At 35 °C (**Figure 17C, l**ower panel), genotypes show more varied responses to heat stress, with some maintaining robust growth while others exhibit reduced height accumulation.

Many of the populations evaluated here are wild and open pollinated, therefore uniform performance among individuals is not expected. The observed within-population variation in GEAVs and phenotypic responses likely reflects this and the differing degrees of relatedness among individuals from the same population. The extent of inbreeding within these populations remains unknown and may influence the ease with which genomic predictions can be directly validated, particularly if there are limited numbers of accessions per population. The present validation focused exclusively on seedling stage responses to temperature dress, and while early growth responses are informative, extending this analysis to later developmental stages and other relevant traits will be necessary to fully assess the predictive utility of Bio 5 and indeed other climate associated GEAVs. Together these results indicate that GEAVs for bioclimatic traits can capture biologically relevant variation in population responses, even within genetically heterogeneous open pollinated populations. Although performance varied among individuals, populations exhibiting consistently elevated Bio 5 GEAVs tended to show more favourable phenotypic responses under elevated temperatures (**Figure S17C**). This suggests that climate associated genomic predictions may aid in pre-breeding by identifying populations enriched for adaptive variation relevant to future climate warming scenarios.

### Genome Environment Associations underlying Climate Adaptation

Loci identified by GEA analysis across temperature related bioclimatic variables (Bio 1 - 11, **Figure 4A, Table S12**) and replicated across multiple variables (≥ 3) were enriched for genes involved in transcriptional regulation, protein homeostasis and RNA metabolism (**Table S13**). Notable examples include histone acetyltransferase MCC1 (NC_044371.1, position 38736665, XP 030489336.1), transcriptional corepressor LEUNIG (NC_044370.1, position 104264844, XP 030485585.1), bHLH3 transcription factor (NC_044375.1, position 5315096, XP 030504442.1) and lysine specific demethylase JMJ25 (NC_044375.1, position 15077356, XM 030648768.1), highlighting regulatory and epigenetic control as an important component of temperate associated adaptation. Several replicated loci also encoded proteins involved in RNA processing, including tubulin folding cofactor E (NC_044371.1, position 82016675, XM 030633544.1), multiple DEAD-box ATP dependent RNA helicases (NC_044379.1, position 41133006, XM 030628330.1; NC_044377.1, position 49103907, XM 030621901.1) and dicer homolog 2 (NC_044375.1, position 21340029, XM 030650266.1) (**Table S13**). Together, these findings suggest that adaptation to temperature variation in Cannabis is mediated through maintenance of transcriptional and post transcriptional stability, enabling cellular function and resilience under variable thermal conditions.

Across precipitation related bioclimatic variables (Bio 12 - 19, **Figure 4B, Table S14**) multiple replicated loci (≥ 3) were associated with genes involved in membrane transport and osmotic regulation (**Table S15**). These included UDP-glycosyltransferase 92A1 (NC_044370.1, position 7720425, XP 030487896.1), a mitochondrial outer membrane protein porin (NC_044372.1, position 11784825, XP 030492781.1), a mitochondrial alkaline neutral invertase (NC_044376.1, position 4400010, XM 030652883.1) and a pyrophosphate-energized vacuolar membrane proton pump (NC_044377.1, position 9932245, XP 030510341.1) (**Table S15**). Collectively, these results highlight the importance of genes regulating intracellular solute flux and proton gradients, key processes involved in cellular homeostasis, maintaining plant performance under variable precipitation conditions.

### Implications for Conservation for abiotic stress tolerance

Globally, species distribution models (SDM) project declines in climatically suitability areas for *Cannabis* (Halpin-McCormick et al., 2025). More specifically, in the Midwest, suitability loss is predicted in areas of Kansas, Illinois, Missouri, Indiana and South Dakota, with expansion of suitable area observed for Iowa and Ohio (Ford et al., 2024). The Cannabis populations sampled in this instance from North America exist in relatively few climate classes, predominantly Dfa (Cold, no dry season, hot summer) in the present day (**Figure 3B, Figure S6**). However, if statewide climate class representation is taken into account, such as those states sampled in the Aina dataset (North Dakota, South Dakota, Nebraska, Kansas, Iowa, Minnesota, Wisconsin, Illinois, Indiana, New York), exhibit substantial heterogeneity in the extent of Köppen-Geiger climate class change from 1901 to the present, with further divergence projected under future climate scenarios (2041-2099) (**Figure S18**). Several Midwestern and Northeastern states exhibit particularly pronounced shifts in dominant climate class through time (**Figure S18A**). For example, the states of Kansas and Indiana show transitions from a cold climate class (25 - Dfa) toward a temperate climate class (14 - Cfa), with New York also exhibiting comparable reduction in cold climate class (26 - Dfb), and shifting from warm to hot summers (25 - Dfa) and later to a temperate climate class (14 - Cfa) (**Figure S18A**). These directional shifts are also evident when examining the proportion of state areas undergoing Köppen-Geiger class transitions between consecutive time points (**Figure S18B**).

State level climate transitions may provide important context for interpreting population specific genomic signals in the Aina dataset. Kansas populations, consistently rank among the top GEAV performers for Bio 5 (maximum temperature of the warmest month) (**Figure S17A**) with the state experiencing an increase from cold (25 - Dfa (1961-1990)) to a warm temperate climate class (14 - Cfa, (1991-2020, 2041-2070)) and may explain the uniformly high Bio 5 GEAVs observed across Kansas populations (**Figure S17A, Figure S16A**). In field trials however, variable performance for Kansas samplings was observed (**Figure S17B**) and in a controlled environment experiment (**Figure S17C**). Indiana and New York populations originate from regions characterised by cold climate classes (25 - Dfa and 26 - Dfb respectively) and may explain why the New York population exhibited one of the lowest population level Bio 5 GEAVs (**Figure S17A**) and reduced performance in field trials (**Figure S17B**). However, interpretation of population level responses is complicated by uncertainty surrounding the genetic stability and breeding history of sampled accessions, which may contribute additional variance to both genomic predictions and phenotypic performance. When considering vulnerable populations, states projected to experience high proportions of Köppen-Geiger climate class change (**Figure S18B**) may face increased risk, as rapid or extensive climatic turnover may outpace local adaptive capacity (Jump & Peñuelas, 2005, Wilczek et al., 2014). Broader sampling would both mitigate current sampling limitations and provide critical insight into adaptive variation that may be at risk of erosion or loss.

### Implications for Breeding for future climate

Future breeding will rely on diverse genetic resources in order to combat changing climates as well as increasing biotic pressures. Having partially validated GEAVs for the populations creates an opportunity for selecting populations to help in recurrent selection breeding programs for climate resilience. This information could be expanded and used by active breeding programs within countries or regions to preserve local landrace material as a future investment for allelic mining as breeding programs develop. The USDA ARS Hemp germplasm repository is an excellent resource and continued characterization of collections like this will provide breeders with long term security of genetic resources. Collections with curated passport records including geographic origins allow for more targeted choice of parents for both allele mining or population improvement. Further characterization of existing landraces will make more clear what populations may harbour beneficial traits. It is very likely that additional populations in North America possess strong associations with their local environment and climate niches, representing an unexplored source of adaptive variation. Continued systematic phenotypic and genomic evaluation of these feral populations in North America will be an important next step for translating this diversity into applied breeding programs.

## Acknowledgements

We would like to thank the University of Hawai’i High Performance Compute Cluster - Koa.

## Author Contributions

Anna Halpin-McCormick - Conceptualization, Formal analysis, Writing, Revising. Ademola Aina - Formal analysis, Writing, Revising.

Michael B. Kantar - Conceptualization, Writing, Revising. Shelby Ellison - Conceptualization, Writing, Revising.

## Conflict of Interest

The authors declare that there is no conflict of interest regarding the publication of this article.

## Data availability

Data and scripts are available at https://github.com/ahmccormick/Feral_Cannabis and at Figshare https://figshare.com/authors/Anna_H_McCormick/17741367

## Tables

**Table S1**. Sample identifiers and geographic coordinates for 909 accessions from the combined Ren et al. (2021), Soorni et al. (2017), and Aina et al. (2025) datasets.

**Table S2**. Summary of the 191 samples with unique latitude longitude coordinates used as the training population across the combined datasets, including associated Köppen-Geiger climate classifications for present and projected future conditions

**Table S3**. Prediction accuracies (mean ± SD) for environmental genomic prediction (EGS) across 19 bioclimatic variables using four genomic prediction models in the combined dataset (n = 909).

**Table S4**. GEAVs for 909 samples across the 19 WorldClim bioclimatic variables.

**Table S5**. Sample identifiers and geographic coordinates for 760 samples from the Aina et al., 2025 dataset.

**Table S6**. Prediction accuracies (mean ± SD) for environmental genomic prediction (EGS) across 19 bioclimatic variables using four genomic prediction models for the Aina dataset (n = 760).

**Table S7**. GEAVs for 760 samples across the 19 WorldClim bioclimatic variables.

**Table S8**. Cannabinoid phenotype measurements for 276 accessions from the Aina dataset.

**Table S9**. Prediction accuracies (mean ± SD) for four cannabinoid traits.

**Table S10**. GEBVs for 760 samples for cannabinoid traits.

**Table S11**. Functional annotations for the top 1% of SNPs by absolute marker effect size contributing to CBD genomic estimated breeding values.

**Table S12**. Functional annotations of SNPs exceeding the WZA significance threshold for temperature related bioclimatic variables (Bio 1 - Bio 11).

**Table S13**. Loci that exceeded the WZA significance threshold (−log10(p) ≥ 2) in three or more temperature-related WorldClim variables (Bio 1–Bio 11) and their annotations.

**Table S14**. Functional annotations of SNPs exceeding the WZA significance threshold for precipitation related bioclimatic variables (Bio 12 -Bio 19).

**Table S15**. Loci that exceeded the WZA significance threshold (−log10(p) ≥ 2) in three or more precipitation-related WorldClim variables (Bio 12 - Bio 19) and their annotations.

**Table S16**. Representative genotypes with the highest and lowest GEAVs for Bio 5 selected for the growth chamber experiment.

**Figure S1.**
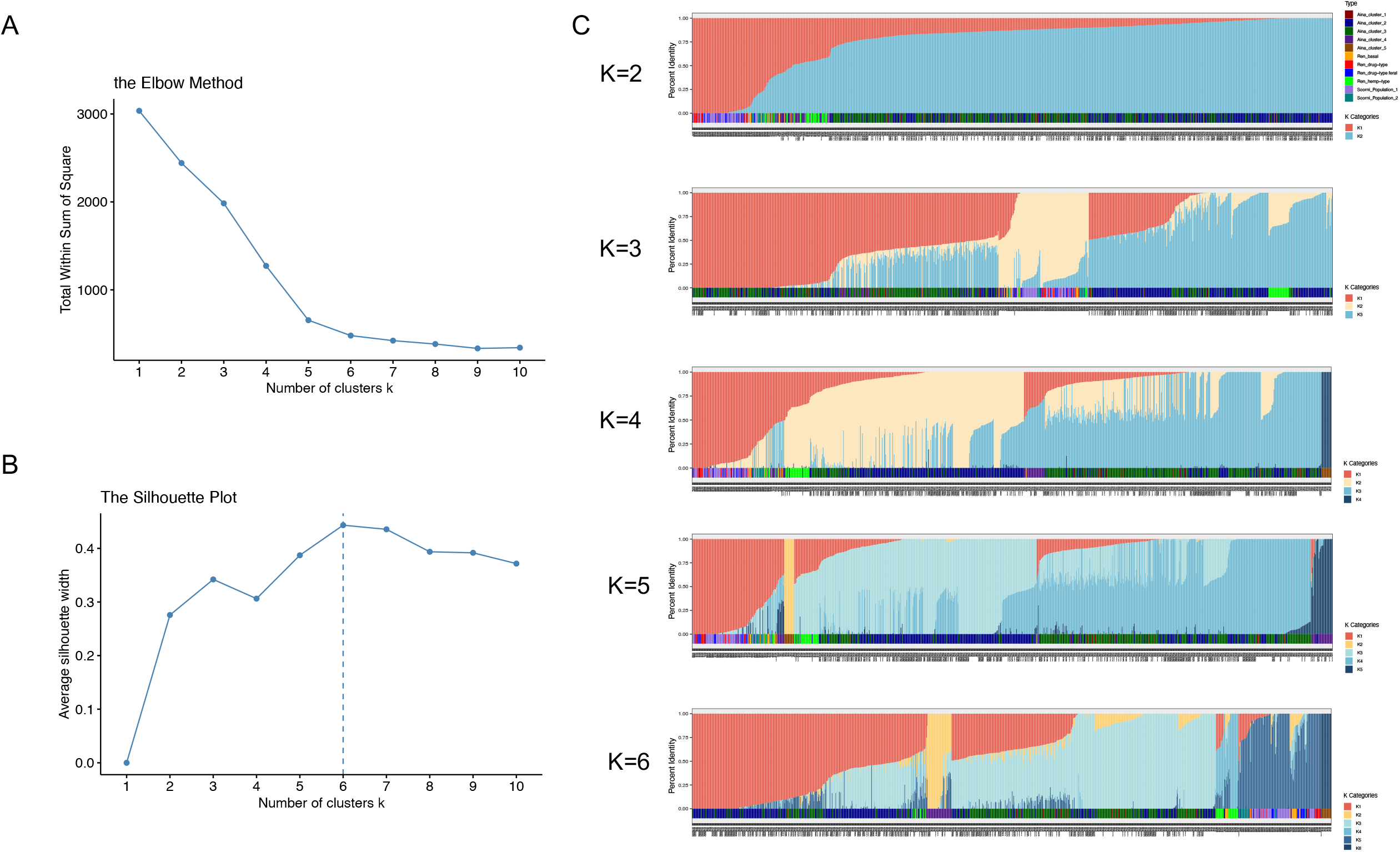
Determination of the optimal number of genetic clusters (K) and population structure inferred using fastSTRUCTURE (**A**) Elbow method showing for K = 1-10 (**B**) Silhouette method showing for K = 1-10 with maximum silhouette score observed at K = 6 (dashed line) (**C**) fastSTRUCTURE bar plots illustrating ancestry proportions for K = 2-6. Each vertical bar represents an individual (n = 909) with colours indicating the proportional assignment to each inferred genetic cluster. Analyses were performed using 13,389 SNPs.

**Figure S2.**
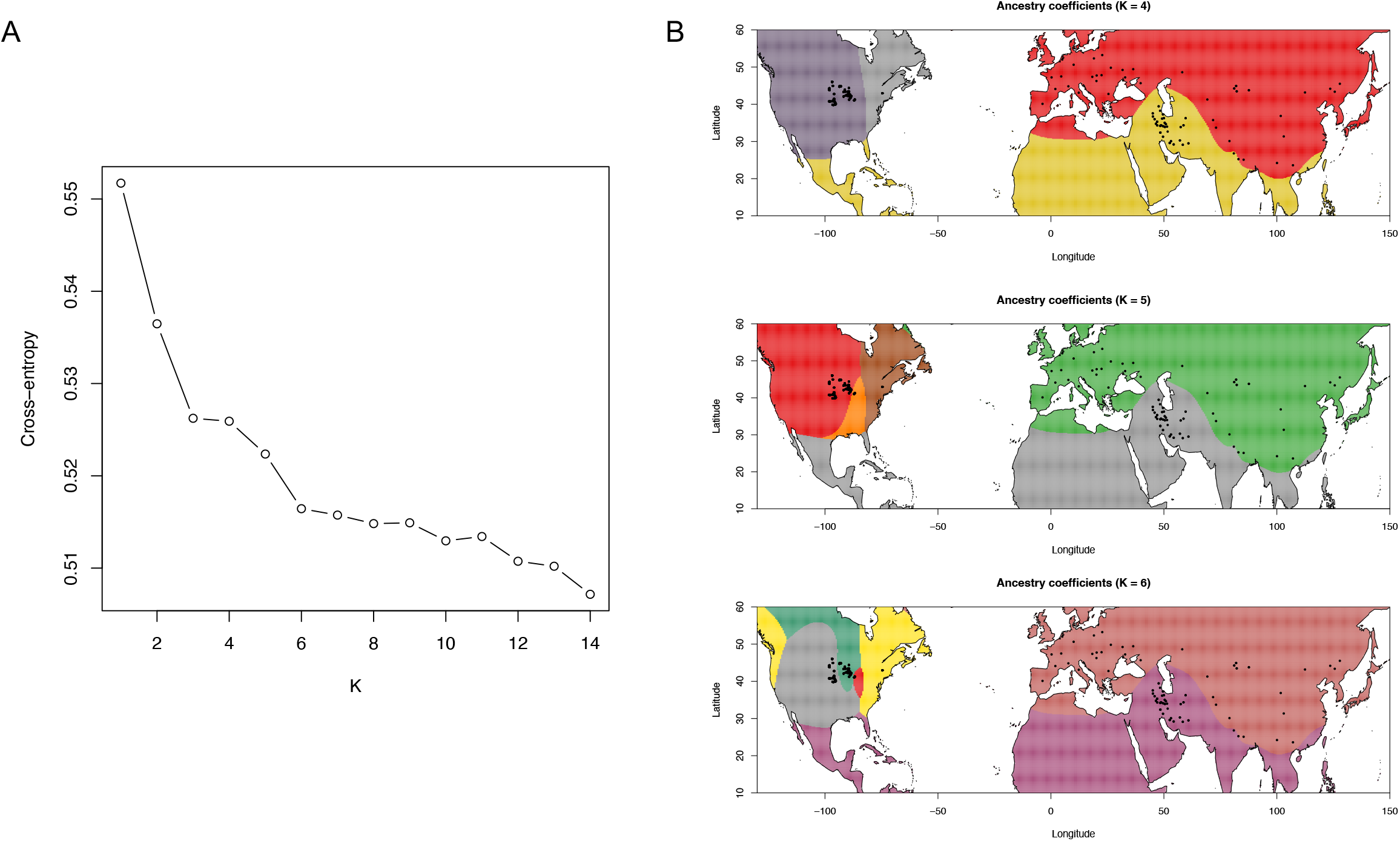
Determination of the optimal number of ancestral populations (K) using TESS3 (**A**) Cross-entropy criterion across values of K (K = 1-14), with decreasing cross-entropy indicating improved model fit (**B**) Spatial ancestry coefficient maps inferred using TESS3 for K = 4, K = 5 and K = 6. Colours represent the inferred ancestry proportions for each genetic cluster, interpolated across geographic space. Black points indicate sample locations. Analysis was performed on 909 samples using 13,389 SNPs.

**Figure S3.**
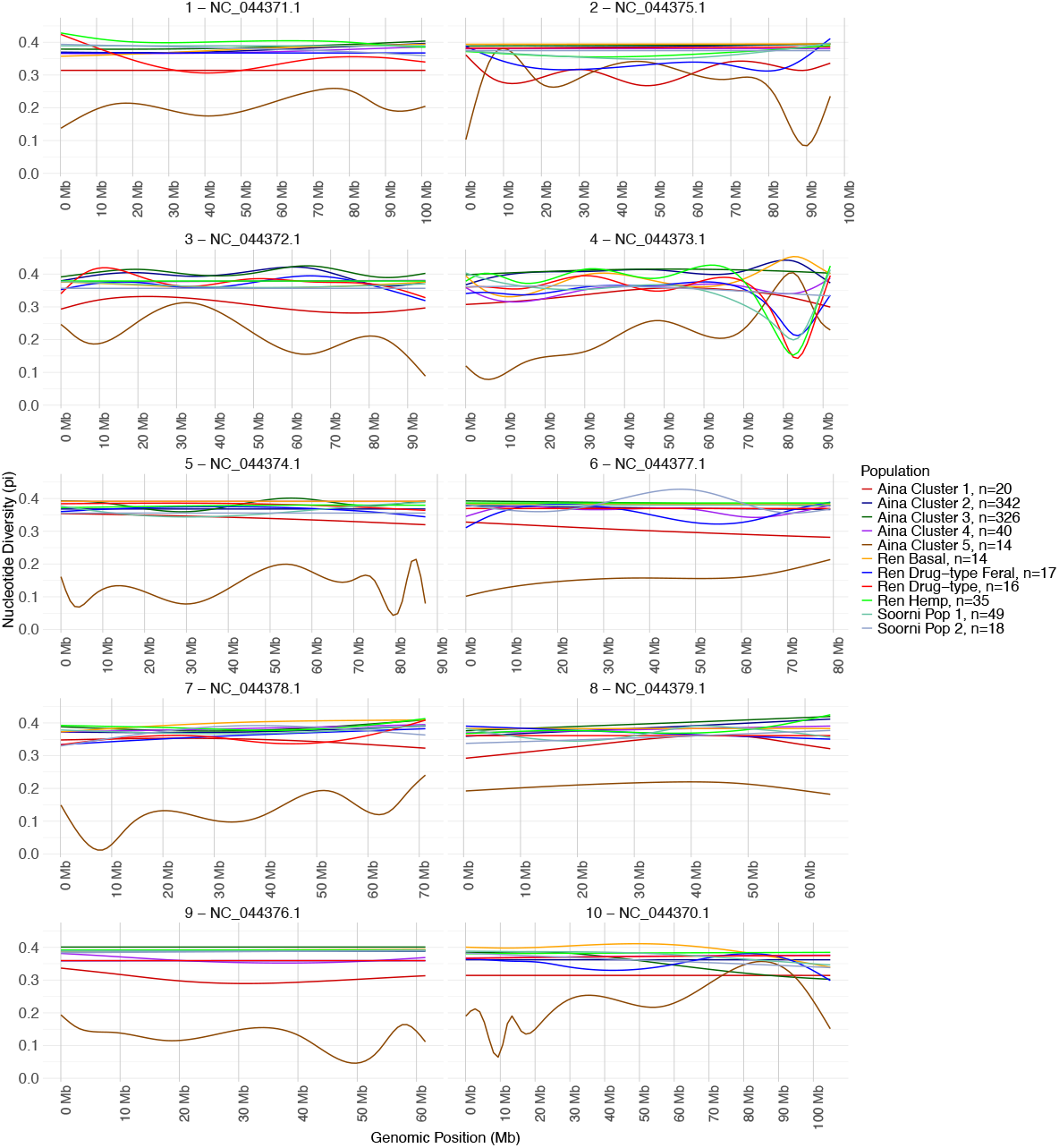
Population-level nucleotide diversity (π) estimated across the genome.

**Figure S4.**
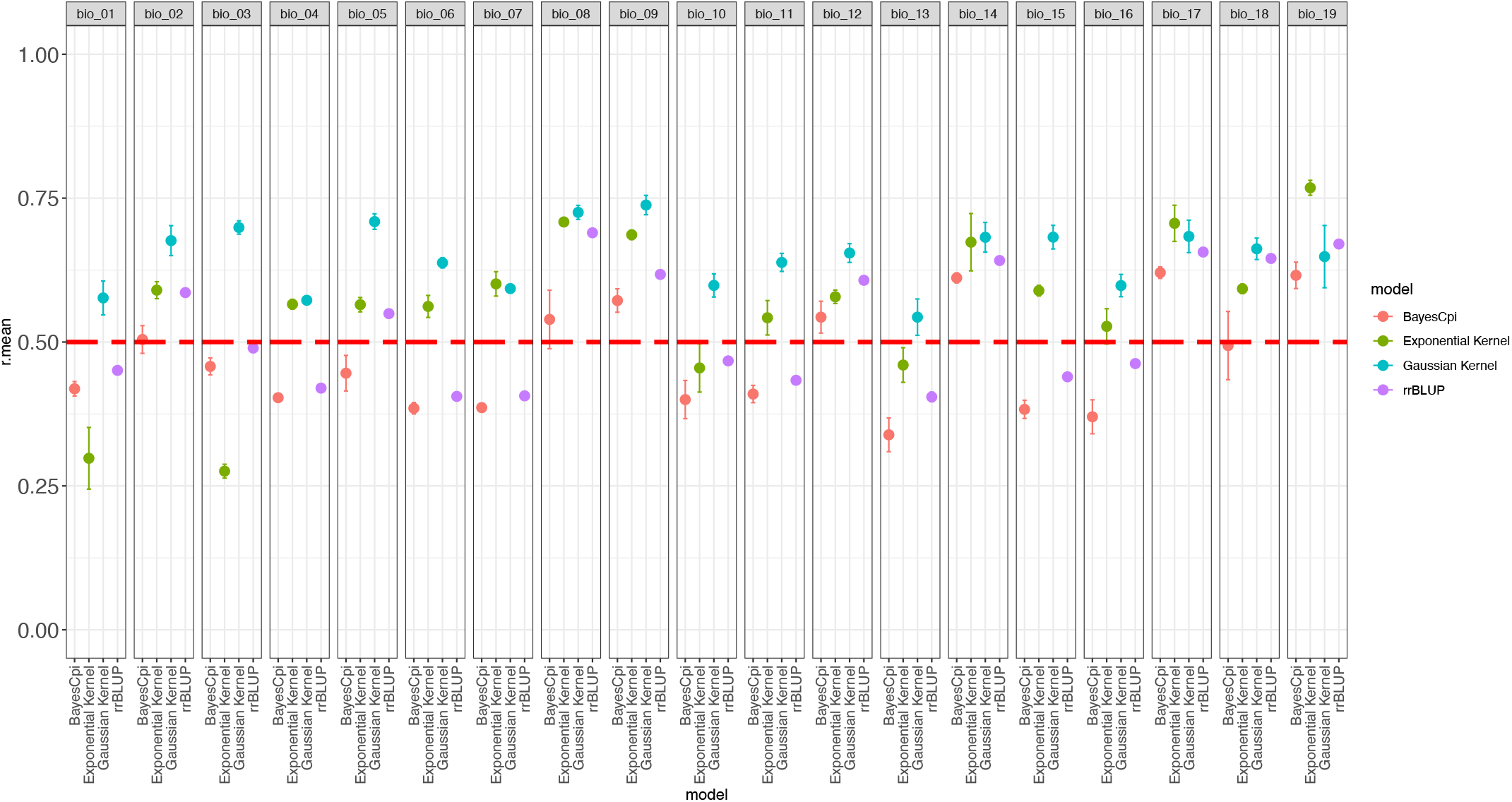
Prediction accuracies for environmental genomic selection (EGS). Models were trained on 191 samples with latitude and longitude information, comprising 80 samples from the Aina dataset, 44 from the Ren dataset and 67 from the Soorni dataset and evaluated on the remaining 718 samples (total n = 909). The original dataset contained 13,389 SNPs, which was filtered to remove loci with >20% missing data, resulting in 12,030 SNPs. Residual missingness (6.19%) was imputed using marker mean imputation. The red dashed line indicates a prediction accuracy threshold of 0.5.

**Figure S5.**
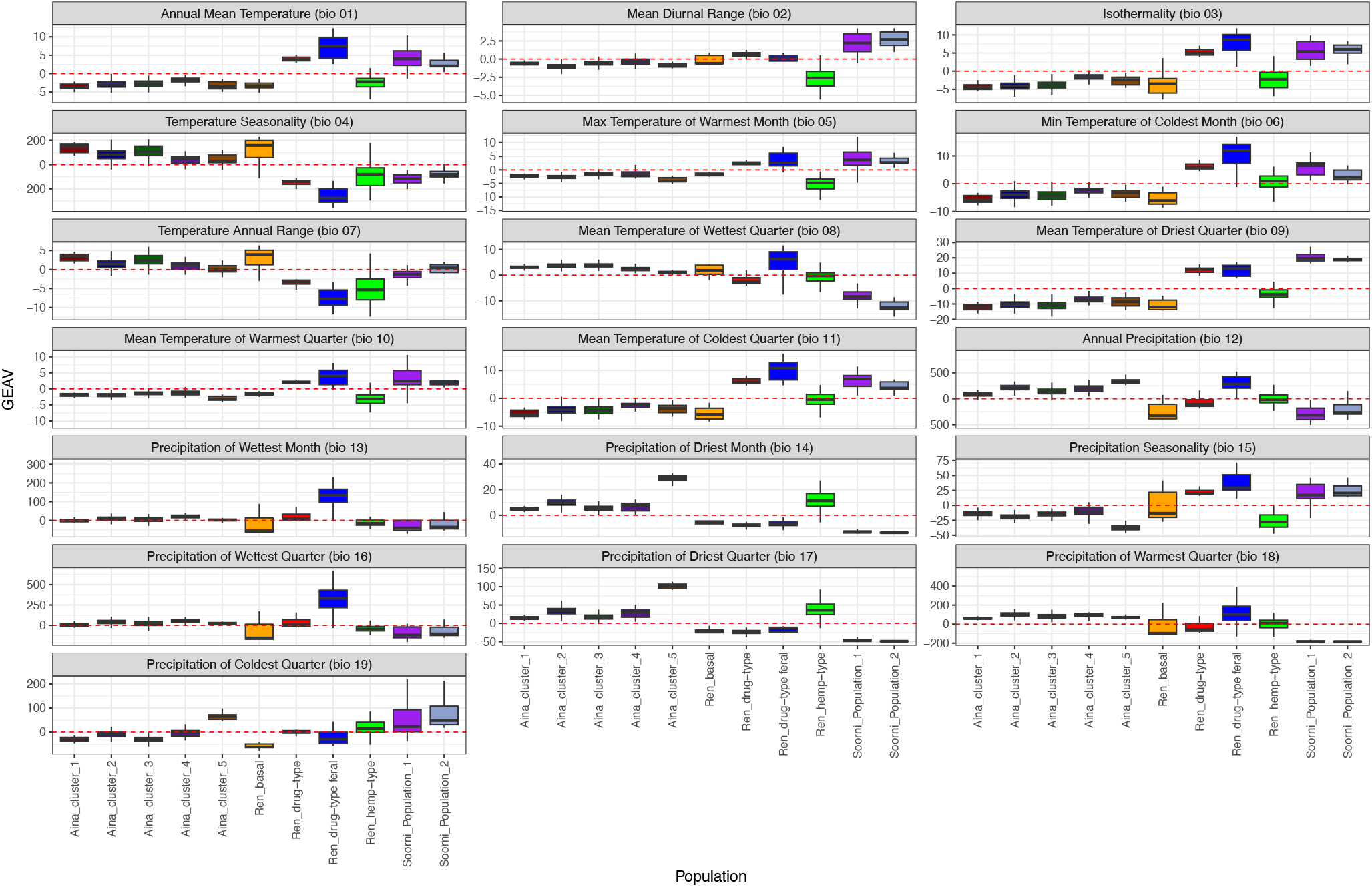
EGS on Eurasian and North American Cannabis datasets (n=909) using 12,959 SNPs for the 11 population groupings for all 19 WorldClim bioclimatic traits.

**Figure S6.**
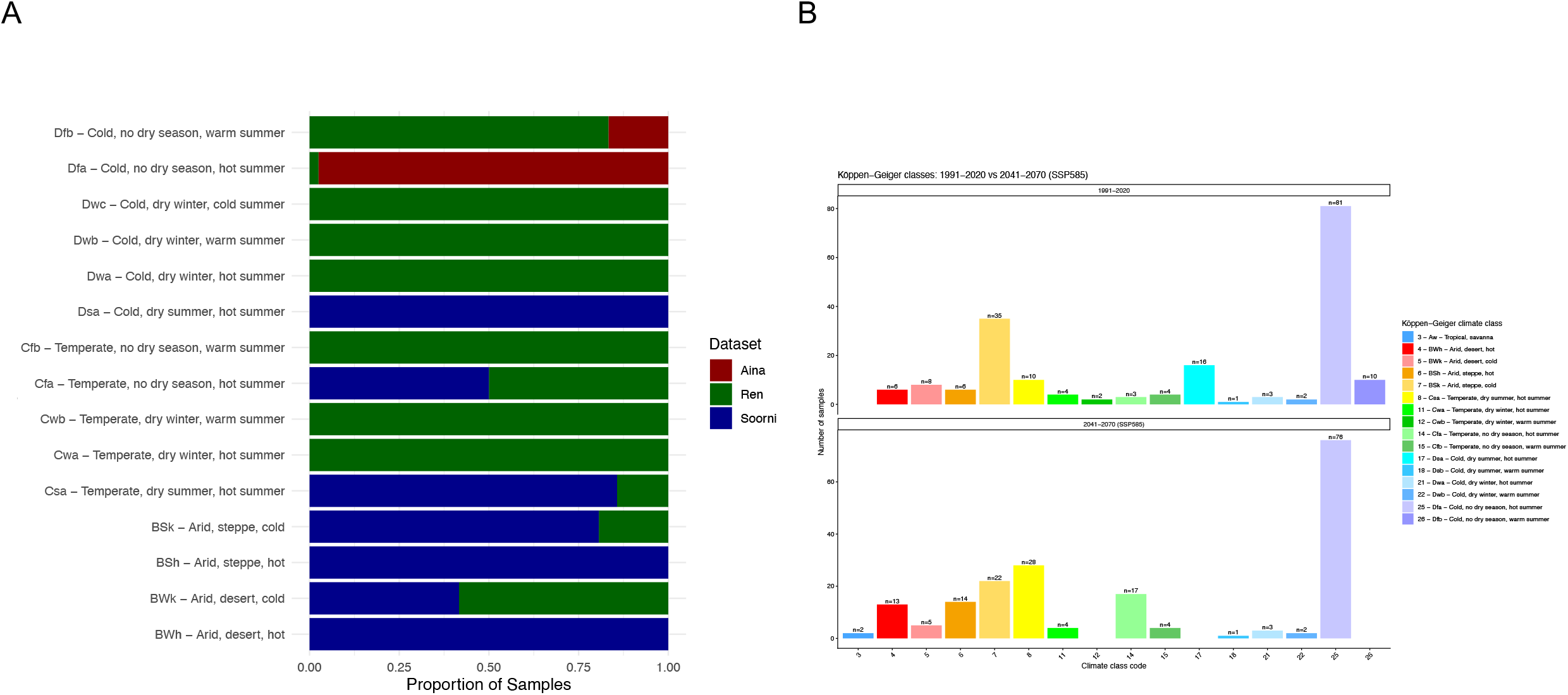
Köppen-Geiger climate classes represented by 191 unique latitude and longitude positions (**A**) By dataset (**B**) Changes in Köppen-Geiger climate classes for all 191 samples with latitude and longitude values from (1991-2000) to (2041-2070) for SSP585.

**Figure S7.**
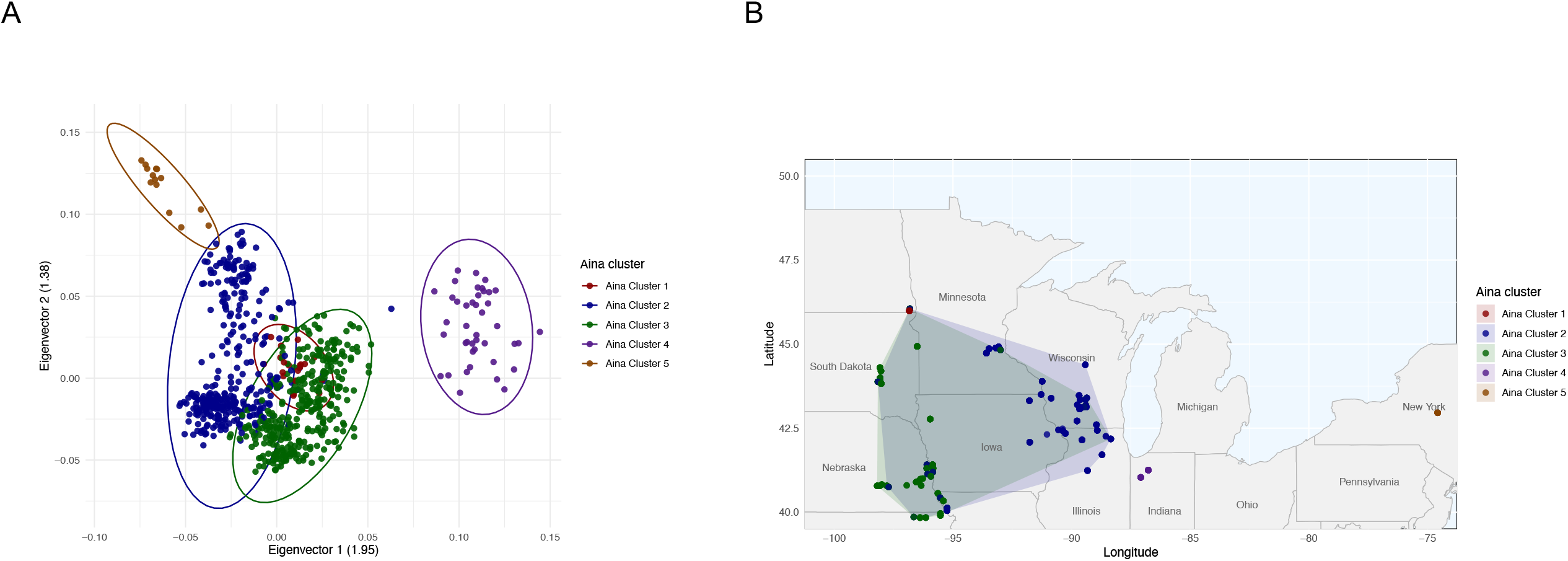
Population structure and geographic clustering of feral *Cannabis* samples (**A**) PCA of 760 feral hemp samples with 3,762 SNPs (LD=0.2) (**B**) Geographic distribution of 742 individual feral samples across 85 population level collection sites. Datapoints are coloured by their hierarchical clustering of principal component (HCPC) cluster assignment. The number of individual samples associated with each cluster is indicated in figure legend.

**Figure S8.**
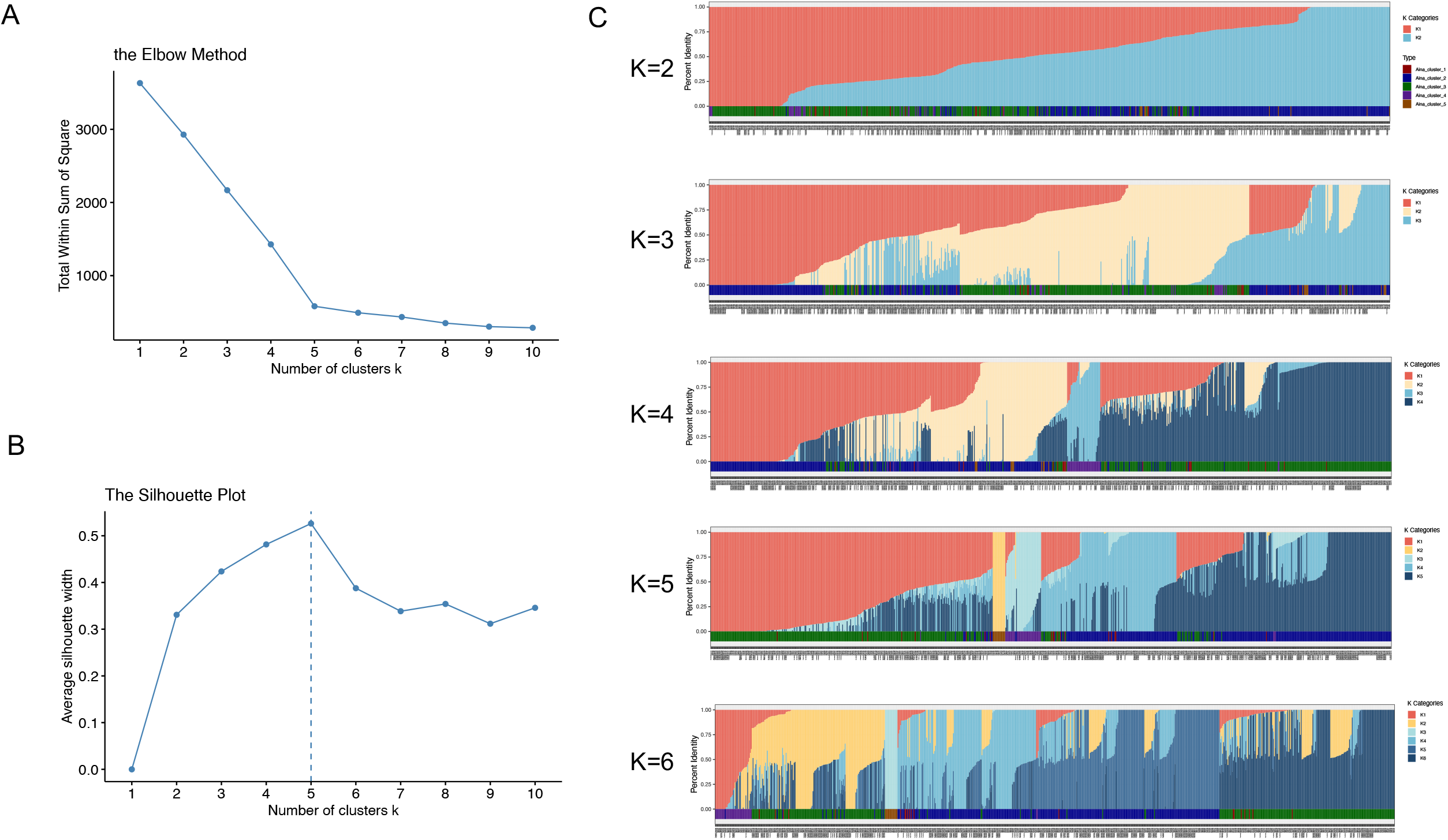
Determination of the optimal number of genetic clusters (K) and population structure inferred using fastSTRUCTURE for the Aina dataset (n=760) (**A**) Elbow method showing for K = 1- 10 (**B**) Silhouette method showing for K = 1-10 with maximum silhouette score observed at K = 5 (dashed line) (**C**) fastSTRUCTURE bar plots illustrating ancestry proportions for K = 2-6. Each vertical bar represents an individual (n = 760) with colours indicating the proportional assignment to each inferred genetic cluster. Analyses were performed using 37,346 SNPs.

**Figure S9.**
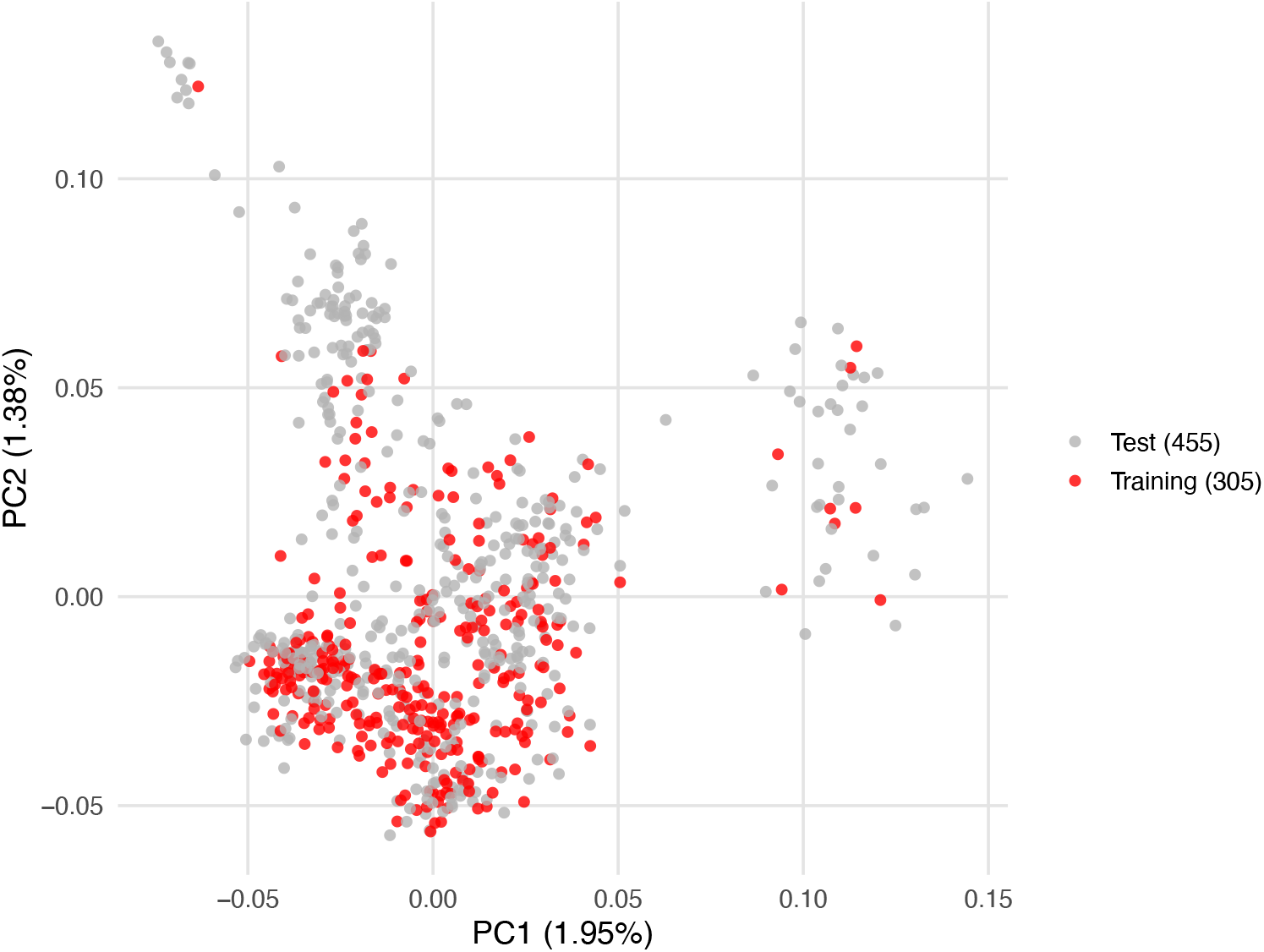
Distribution of training lines (n=305) used for EGS with representation in each HCPC cluster. The training set of 305 samples comprised 3/20 samples from cluster one, 133/342 from cluster 2, 161/326 from cluster 3, 7/40 from cluster 4 and 1/14 from cluster 5.

**Figure S10.**
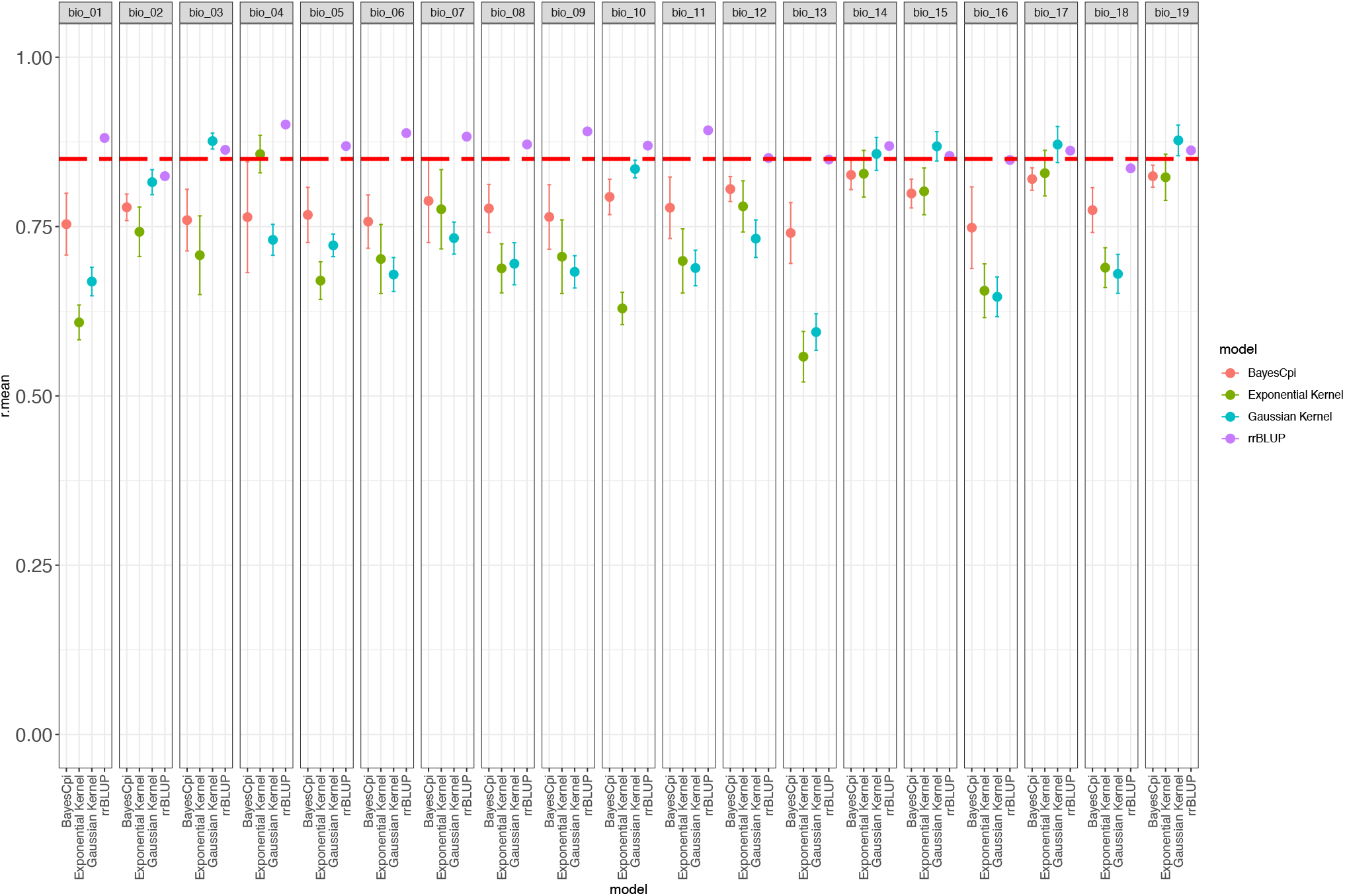
Prediction accuracies for environmental genomic selection (EGS). Models were trained on 305 samples from the Aina. The original dataset contained 37,346 SNPs, which was filtered to remove loci with >20% missing data, resulting in 26,433 SNPs. Residual missingness (8.3%) was imputed using marker mean imputation. The red dashed line indicates a prediction accuracy threshold of 0.85.

**Figure S11.**
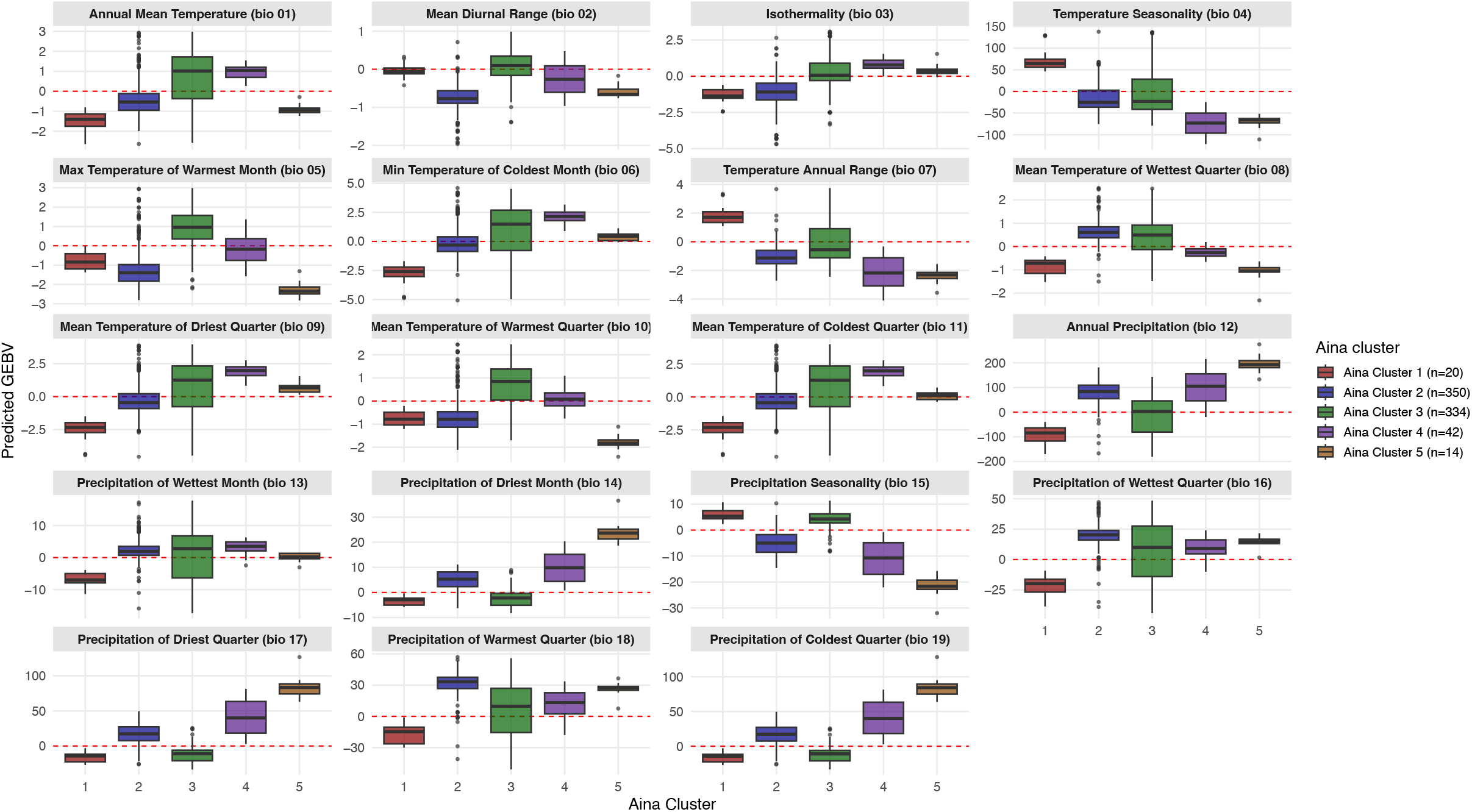
GEAV results by HCPC cluster for the North American (Aina) Cannabis dataset (n=760) for all 19 WorldClim bioclimatic traits with 22,852 SNPs. A training set of 305 (∼40% of the dataset) was developed by maximizing genetic distance for sample selection leaving 455 samples in the test set.

**Figure S12.**
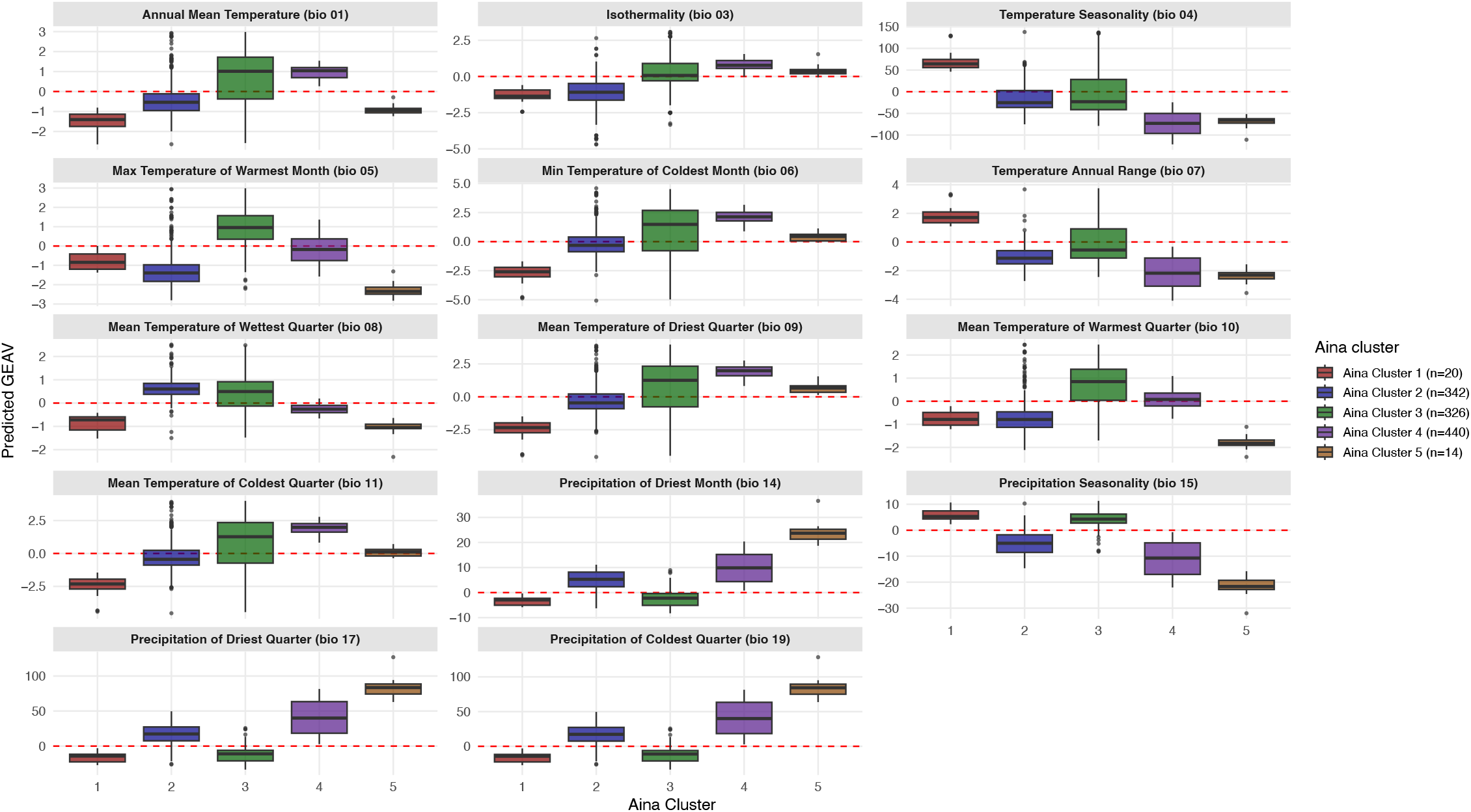
GEAV results by HCPC cluster for the North American (Aina) Cannabis dataset (n=760) with 22,852 SNPs. Fourteen WorldClim bioclimatic traits had prediction accuracies greater than 0.85

**Figure S13.**
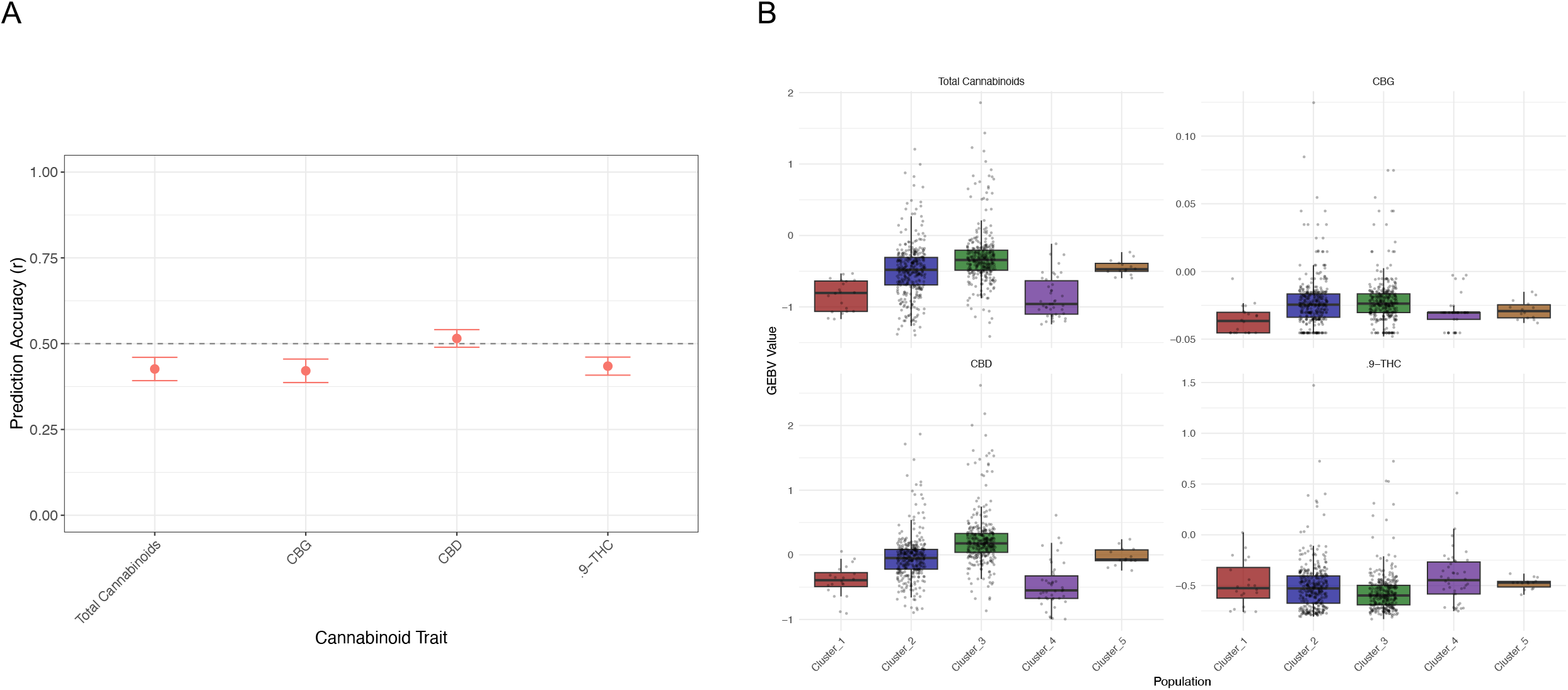
Prediction accuracies (PAs) and genomic estimated breeding values (GEBVs) for four cannabinoid traits by population (**A**) Prediction accuracies (PAs) and (**B**) genomic estimated breeding values (GEBVs) for four cannabinoid traits, summarized by population. Genomic selection models were trained on 273 samples and evaluated on 487 test samples. A total of 37,346 SNPs were used for genomic selection, with 14.65% missing data imputed using marker mean imputation.

**Figure 14.**
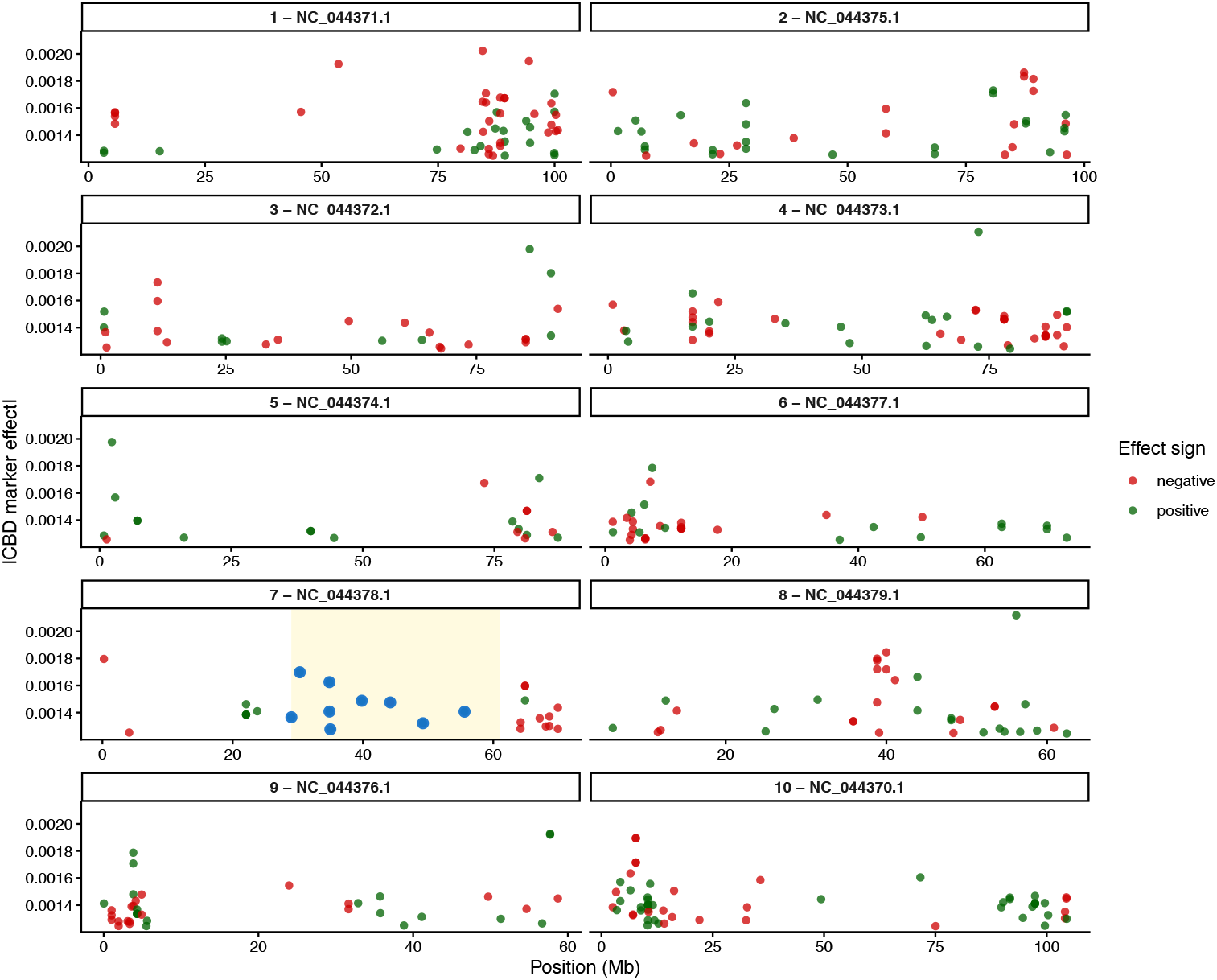
Genomic distribution of large effect markers (top 1% by absolute rrBLUP effect size, n=374) for CBD concentration. 9 SNPs in the CBDAS cluster (29,635,083 – 61,400,085) on chromosome 7 are highlighted in the yellow box.

**Figure S15.**
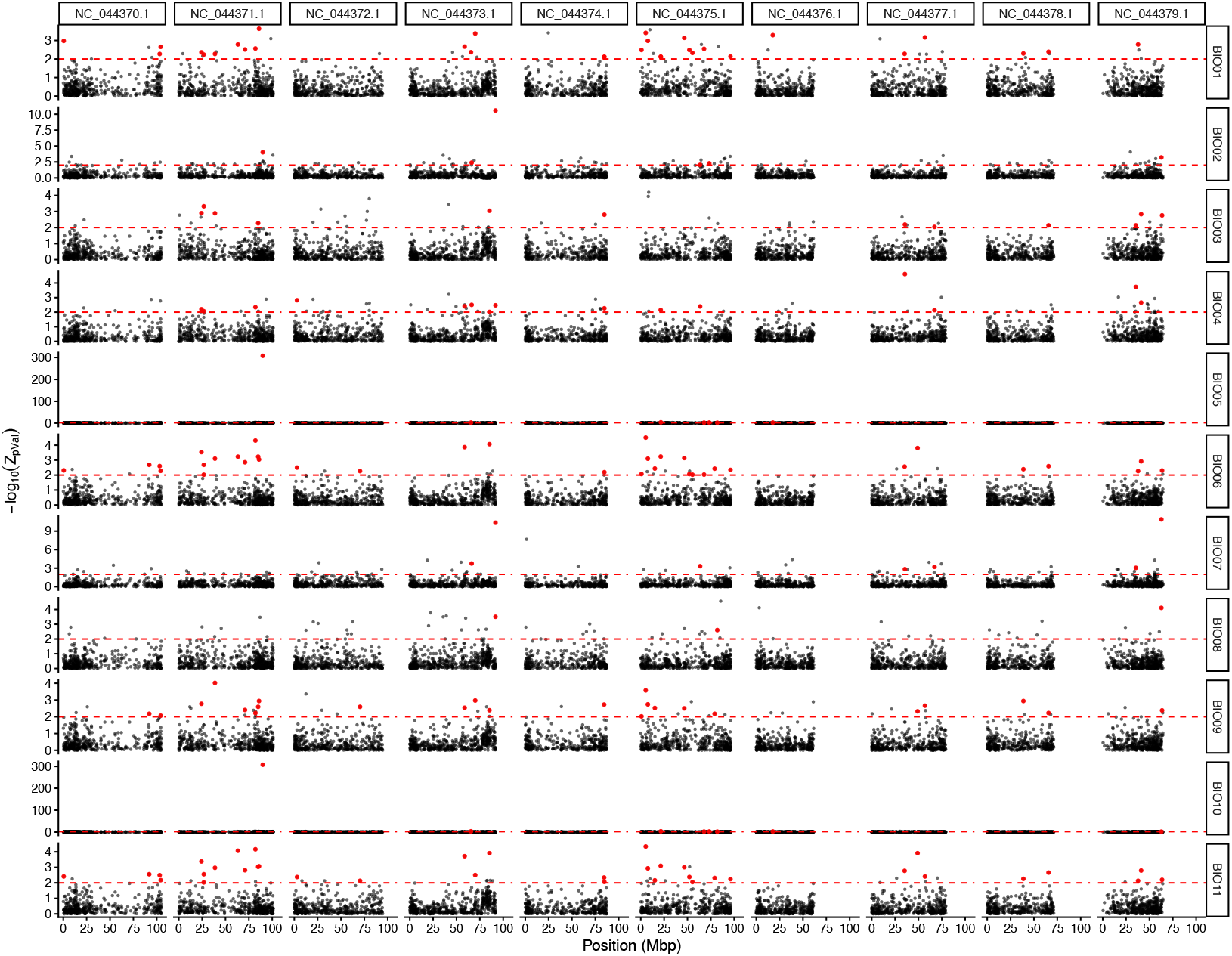
Genome-environment associations using Kendall’s τ between population allele frequency for the WorldClim temperature related variables (Bio 1 - 11) across 10 populations. SNP-level p-values were aggregated into 50-kb window scores using the Weighted Z-analysis (WZA). The red dashed line represents -log10(0.01). Red points indicate loci that exceeded the WZA significance threshold in at least three temperature-related WorldClim variables (Bio 1 - Bio 11).

**Figure S16.**
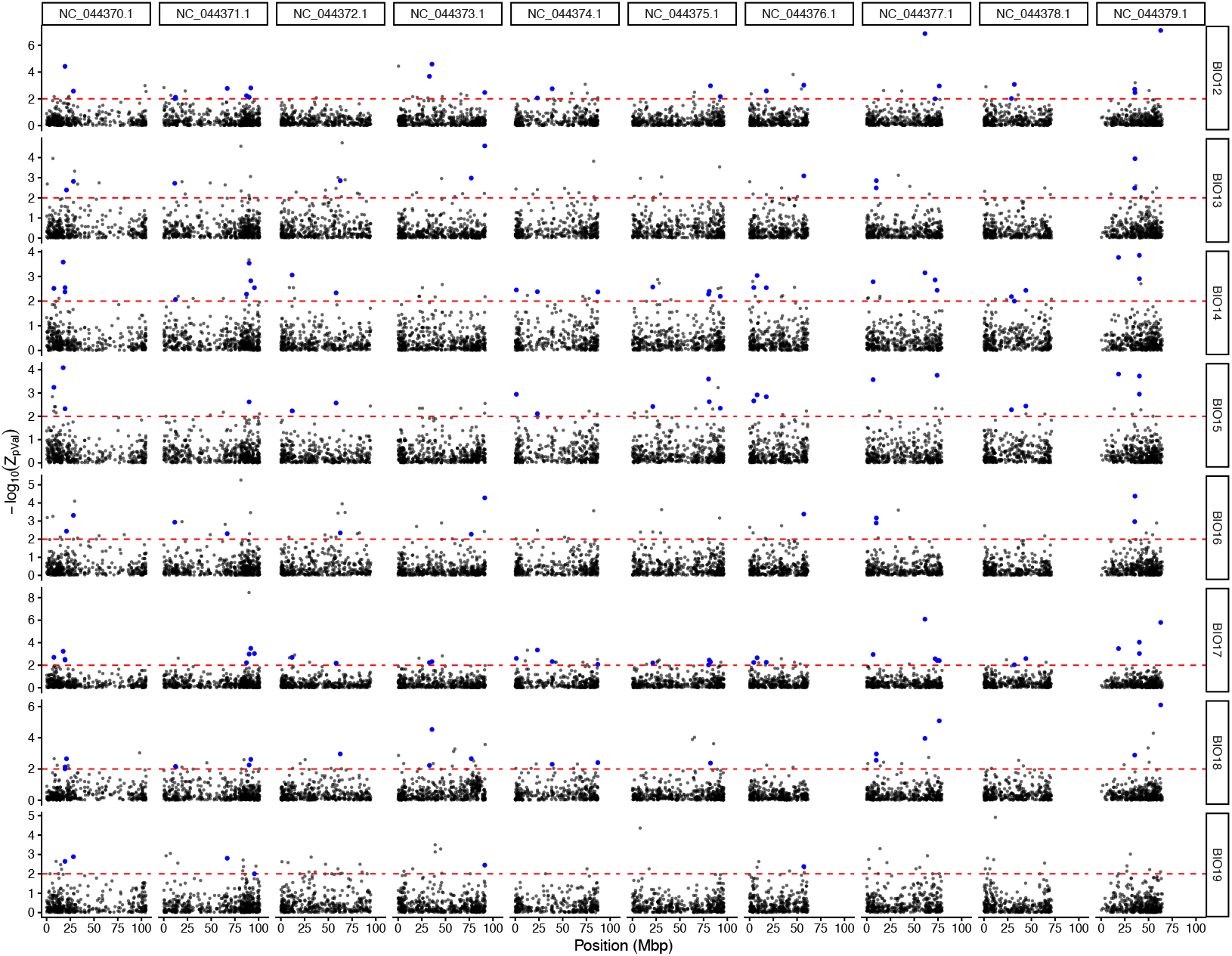
Genome-environment associations using Kendall’s τ between population allele frequency for the WorldClim precipitation related variables (Bio 12 - 19) across 10 populations. SNP-level p-values were aggregated into 50-kb window scores using the Weighted Z-analysis (WZA). The red dashed line represents -log10(0.01). Blue points indicate loci that exceeded the WZA significance threshold in at least three precipitation-related WorldClim variables (Bio 12 - Bio 19).

**Figure S17.**
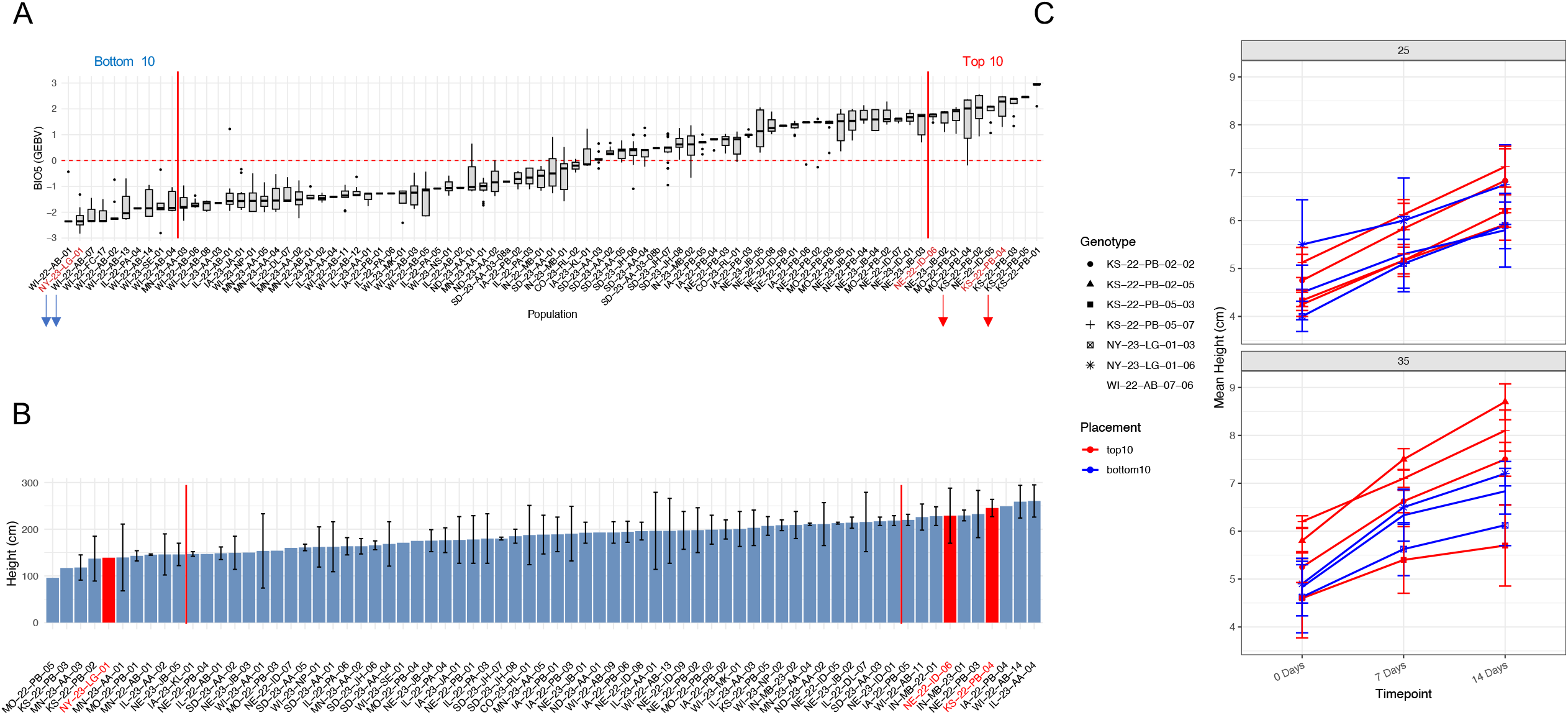
Genomic temperature associations and heat-dependent growth variation (**A**) Population level (n=85) GEAVs for Bio 5 (max temperature of the warmest month) (**B**) Field evaluation of feral hemp populations for plant height (cm). Red dashed line indicates the top and bottom 10 populations. Samples in red represent those which are positioned in the top 10 or bottom 10 GEBVs for Bio 5 (temperature of the warmest month) from the above plot in panel A (**C**) Mean plant height across three developmental timepoints (0, 7, and 14 days) for seven cannabis genotypes grown under two temperature treatments (25°C and 35°C). Lines represent genotype-specific growth trajectories, with error bars indicating standard error. At 25°C (upper panel), genotypes exhibit relatively parallel growth patterns with consistent separation among lines. At 35°C (lower panel), genotypes show more varied responses to heat stress, with some maintaining robust growth while others exhibit reduced height accumulation.

**Figure S18.**
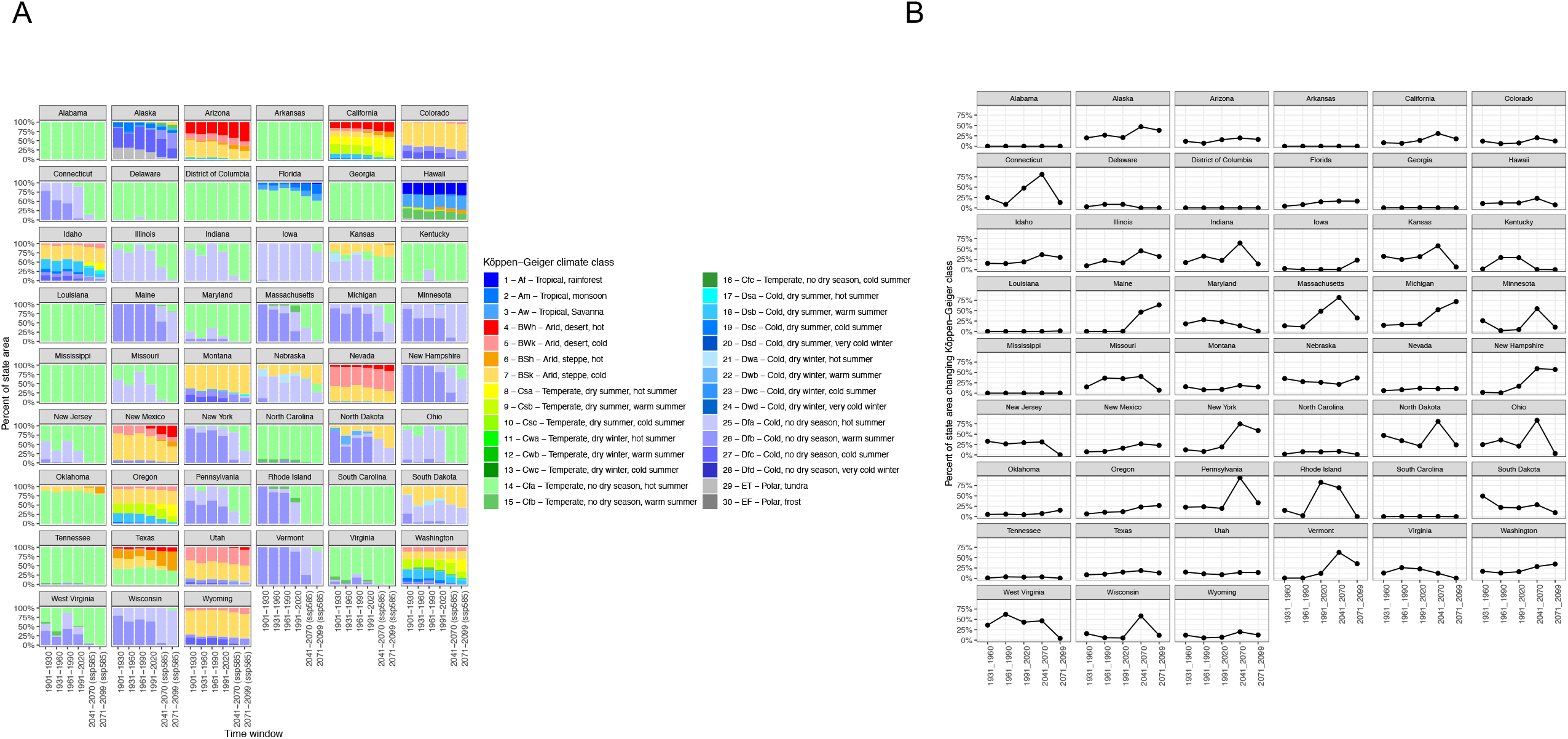
Köppen-Geiger climate composition and turnover through time for the United States (**A**) Changes in proportions of Köppen-Geiger climate classes per state through time (**B**) Percent of state area changing Köppen-Geiger climate class through time. For each state, proportional climate class composition and the fraction of area undergoing class transitions between consecutive periods were calculated from rasters using spatial overlays, where each grid cell represents approximately a 10 × 10 km area and is assigned a single climate class.

## Notes

https://figshare.com/authors/Anna_H_McCormick/17741367

https://github.com/ahmccormick/Feral_Cannabis

### Competing Interest Statement

The authors have declared no competing interest.

